# Haplotype-based insights into the genetic architecture of net blotch resistance in barley

**DOI:** 10.64898/2026.07.23.740211

**Authors:** Dan Liu, Xuechen Zhang, Lisle Snyman, Tara Garrard, Hugh Wallwork, Hari Dadu, Mark Maclean, Jingyang Tong, Chensong Chen, Dilani Jambuthenne Gamaralalage, Sambasivam Periyannan, Lee Hickey, Ben J. Hayes, Eric Dinglasan

## Abstract

Net blotch, caused by *Pyrenophora teres*, is a major constraint to barley production worldwide and occurs as two epidemiologically distinct forms: net form net blotch (NFNB) and spot form net blotch (SFNB). Although numerous resistance loci have been reported in recent years, their genetic relationship remains poorly understood, and the effective deployment of resistance is constrained by the complex genetic architecture of net blotch resistance. In this study, we used a haplotype-based mapping approach to dissect the genetic basis of resistance to NFNB and SFNB in a diverse panel of 950 barley accessions from the Australian Grains Genebank (AGG). Disease responses were evaluated across 13 experiments, and a total of 40 quantitative trait loci (QTL) were identified, including 26 associated with NFNB, 29 with SFNB, and 15 common for both diseases. Most loci co-localized with previously reported QTL, while six putative novel haploblocks highlighted untapped genetic diversity within the AGG collection. Correlation analyses across phenotypic, genetic and haploblock levels revealed a partial but incomplete overlap in resistance mechanisms between NFNB and SFNB. Among the 4,497 haploblocks, approximately 60% of them showed positive local genetic correlations between the two diseases, suggesting shared genomic contributions to resistance. Haplotype composition analysis further identified a resistant haplotype group, mainly comprising accessions of Asian origin, that exhibited high levels of resistance to both forms of net blotch. Through *in silico* haplotype stacking, we demonstrated the cumulative genetic potential achievable by combining favourable haplotypes. When the breeding objective was to improve resistance to both NFNB and SFNB, dual-disease stacking strategies outperformed single-disease approaches, highlighting the value of prioritising haplotypes with positive pleiotropic effects. Overall, this study provides a comprehensive haplotype-level framework for understanding net blotch resistance and delivers practical insights for breeding barley cultivars with durable and broad-spectrum resistance to both NFNB and SFNB.

## 1 Introduction

Barley (*Hordeum vulgare* L.) is one of the most important cereal crops in the world, mainly used for animal feed, malt, and human consumption in Japan, Korea and China [1,2]. However, its production is constantly constrained by numerous diseases. Among these, net blotch is one of the most prevalent foliar fungal diseases, occurring almost in all barley-growing regions worldwide, and causing yield losses typically ranging from 10% to 40%, with even greater losses under conductive conditions [3].

Net blotch has two distinct forms, net form net blotch (NFNB) and spot form net blotch (SFNB), caused by *Pyrenophora teres* f. *teres* (*Ptt*) and *Pyrenophora teres* f. *maculata* (*Ptm*), respectively [4]. Although closely related, *Ptt* and *Ptm* represent genetically separate pathogen populations with different symptoms on infected host plants [5,6]. NFNB produces elongated necrotic lesions with longitudinal and transverse striations [7], while SFNB produces discrete oval to circular lesions surrounded by chlorosis or necrosis [8]. Their contrasting symptoms are underpinned by distinct *in planta* growth behaviors. Microscopy studies showed that *Ptm* grows slowly and initially as a biotroph with intracellular vesicle formation, while *Ptt* rapidly establishes extensive intercellular growth within the mesophyll, causing broader necrosis [9]. However, it can be difficult to distinguish the two forms of net blotch and mixed infections are common in the field [10].

*P*. *teres* has a high evolutionary capacity, driven by both sexual and asexual reproduction [11]. Sexual recombination enables rapid reshuffling of virulence alleles [12], whereas asexual reproduction allows advantageous pathotypes to spread rapidly. Moreover, *Ptt* and *Ptm* can co-exist in the field and the hybridization between the two forms is rare but possible [13–15]. These processes substantially increase pathogen genetic diversity and contribute to the emergence of novel and highly virulent pathotypes. For example, recent surveillance conducted in 2024 in the Yorke Peninsula region of South Australia detected *Ptt* isolates carrying mutations associated with resistance to Group 3, 7 and 11 fungicides [16]. In addition, natural recombination between *Ptt* and *Ptm* in Western Australia generated hybrid isolates with altered virulence profiles and exhibiting resistance to azole fungicides [15]. It thus underscores the need for integrated disease management in which host resistance is the most durable and sustainable strategy [17].

Host resistance to net blotch can be broadly classed into qualitative and quantitative forms. Qualitative resistance is generally controlled by one or a few major-effect genes and may produce strong but isolate-specific responses, often involving effector-triggered immunity [18,19]. In contrast, quantitative resistance is generally mediated through pattern-triggered immunity and involves multiple minor-effect loci that contribute broad-spectrum, race-nonspecific, and often more durable resistance [20,21]. Although *P. teres* is predominantly necrotrophic, it exhibits a brief transient biotrophic phase during early infection [9]. Early infection of *P. teres*–barley is consistent with a classic gene-for-gene model, in which a host resistance gene recognizes a corresponding pathogen avirulence factor and activates defence responses that restrict pathogen spread [22]. However, both *Ptt* and *Ptm* can also engage in inverse gene-for-gene interactions, where necrotrophic effectors exploit host defence pathways to induce programmed cell death and promote susceptibility [3,23,24].

Significant progress has been made in characterizing the genetic basis of resistance to both NFNB and SFNB. Over the past three decades, numerous QTL distributed across all seven barley chromosomes have been identified by QTL mapping and genome-wide association studies (GWAS) [3,25,26]. Although few resistance gene has been cloned, eleven resistance genes (*Rpt1*–*Rpt11*) and several susceptibility loci (*Spt1*–*Spt4*) have been formally designated. A recent comprehensive review positioned 432 NFNB and 131 SFNB QTL onto the Morex V3 physical genome, highlighting both shared and disease-specific genomic regions [26]. These findings highlighted the complex and highly polygenic nature of net blotch resistance.

Only a limited number of studies have analysed NFNB and SFNB jointly, even though increasing evidence indicates that the two diseases share parts of their genetic architecture. For example, five of the reported genes (*Rpt3*, *Rpt4*, *Rpt5*, *Rpt6*, and *Rpt7*) were associated with both diseases [3]; Liu et al. (2026) reported 42 genomic regions associated with multiple diseases, including NFNB, SFNB and leaf scald. Targeting these shared genomic regions in breeding could offer opportunities to improve resistance to multiple diseases, potentially enhancing selection efficiency and facilitating the development of cultivars with broad and durable resistance. Nevertheless, resistance breeding is further complicated by the pronounced genotype by environment interactions, particularly those arising from the high variability of pathogen pathotypes and yearly disease outcomes. Moreover, many reported loci remain difficult to deploy in breeding because they spanned large physical intervals that may complicate the breaking of undesirable linkage drag, showed inconsistent effects across environments, or relied on single SNP associations that are unstable during recombination [27].

Haplotype-based approaches offer a powerful alternative for resolving these limitations. By jointly modelling all SNP markers and grouping them into haploblocks, the localGEBV framework captures the collective effect of linked alleles and provides more accurate estimates of functional genomic segments [28,29] This improves the robust detection of target chromosome segments and produces haploblocks that are more readily tracked and introgressed in breeding programs. Several recent studies have demonstrated the effectiveness of haplotype-based strategies for dissecting disease resistance and agronomic traits across barley, wheat, and chickpea [27,30–36].

In this study, we used a diverse panel of 950 barley accessions from the Australian Grains Genebank (AGG), together with multi-environment disease screening, high-density genotyping, and haplotype-based mapping strategies to investigate the genetic architecture of resistance to NFNB and SFNB. The specific objectives were to: (1) explore global (individual-level) and local (haploblock-level) correlations between NFNB and SFNB; (2) identify genomic regions associated with NFNB and SFNB response; and (3) evaluate the cumulative effects for resistance improvement through *in silico* favourable haplotype stacking for single and dual disease resistance. The outcomes of this study will provide new insights into the relationship between NFNB and SFNB, as well as their shared and specific genomic determinants of resistance, offering practical strategies for stacking broad-spectrum resistance in barley breeding programs.

## 2 Materials and Methods

### 2.1 Plant materials

The study utilized a panel of *n* = 950 barley accessions sourced from AGG. These accessions originated from Asian barley core collection, the Spanish barley core collection and USDA barley imports, representing germplasm from 21 countries worldwide, spanning Africa (*n* = 177), Asia (*n* = 446), Europe (*n* = 181), North America (*n* = 143) and Oceania (*n* = 3). Most entries are six-rowed barley (*n* = 712), whereas *n* = 184 are two-row types. In terms of growth habit, *n* = 742 accessions were classified as spring barley and *n* = 85 as winter barley (Table S1).

### 2.2 Disease evaluation

A total of *n* = 13 experiments were conducted to evaluate NFNB and SFNB resistance during 2023 and 2024 (Table 1). Seven field experiments were carried out for NFNB, while six experiments were conducted for SFNB, comprising four field trials and two seedling tests.

#### Adult plant screening for NFNB and SFNB

Adult plant screening for NFNB and SFNB was conducted out at the Queensland Department of Primary Industries (QDPI) and Agriculture Victoria (AgVic) during 2023–2024.

In QDPI, field nurseries were conducted at the Hermitage Research Facility (HRF), Warwick. Hill plots were established in rows spaced 76 cm apart, with 50 cm between plots within each row. Each test plot was bordered by a row of a susceptible spreader line to ensure uniform disease pressure. Artificial inoculation was performed using freshly infected stubble. For NFNB, five field trials (NB73_QDPI_2023, NB73_QDPI_2024, NB85_QDPI_2023, NB85_QDPI_2024, NB350_QDPI_2024) were conducted using three distinct *Ptt* isolates (NB73, NB85, and NB350) across two years. For SFNB, two trials (SFNB_QDPI_2023 and SFNB_QDPI_20244) were performed using the *Ptm* isolate SFNB320.

At AgVic, adult plant screening was carried out at Horsham SmartFarm. Approximately 10 seeds of each barley line were sown as a single row/plot of ∼0.55 m in length, with susceptible spreaders (*cv.* VB9613 for NFNB and *cv.* Dash for SFNB) sown around the outside to facilitate infection. Stubble residues infected with the target disease were evenly applied across the screened line to initiate infection. This resulted in four additional datasets shown in Table 1.

At both locations, disease severity was visually scored on adult plants under field conditions using the 1–9 disease rating scale according to National Variety Trials (NVT) (https://nvt.grdc.com.au/), where 1 = resistant; 2 = resistant–moderately resistant; 3 = moderately resistant; 4 = moderately resistant–moderately susceptible; 5 = moderately susceptible; 6 = moderately susceptible–susceptible; 7 = susceptible; 8 = susceptible–very susceptible; 9 = very susceptible.

#### Seedling test for SFNB

Seedling response to SFNB was evaluated at Horsham controlled environment facility following the method described in [37]. Seedling testing consisted of sowing three replicates of each barley accession into pots containing potting mix and fertilizer arranged in a randomized design. Seedlings were grown to the two to three-leaf stage (Z12-13) and inoculated with a mixed spore suspension of individual pure isolates.

In 2023, six *Ptm* isolates (Ptm22-028, 073, 075, 089, 099 and 134) with contrasting virulence profiles were chosen to screen the AGG germplasm. In 2024, a mixture of six additional isolates (Ptm19-197; Ptm23-023; Ptm23-037; Ptm23-084; Ptm23-057; Ptm23-074) was used (Table 1). Inoculated seedlings were incubated in a humidity tent for 24–32 h. Infected seedlings were then transferred to a glasshouse maintained at 20 ± 2 °C with a 12 h light/dark cycle for 6–8 days to allow symptom development. Each barley accession was visually assessed for disease severity on a 1–9 scale as described above.

**Table 1.**
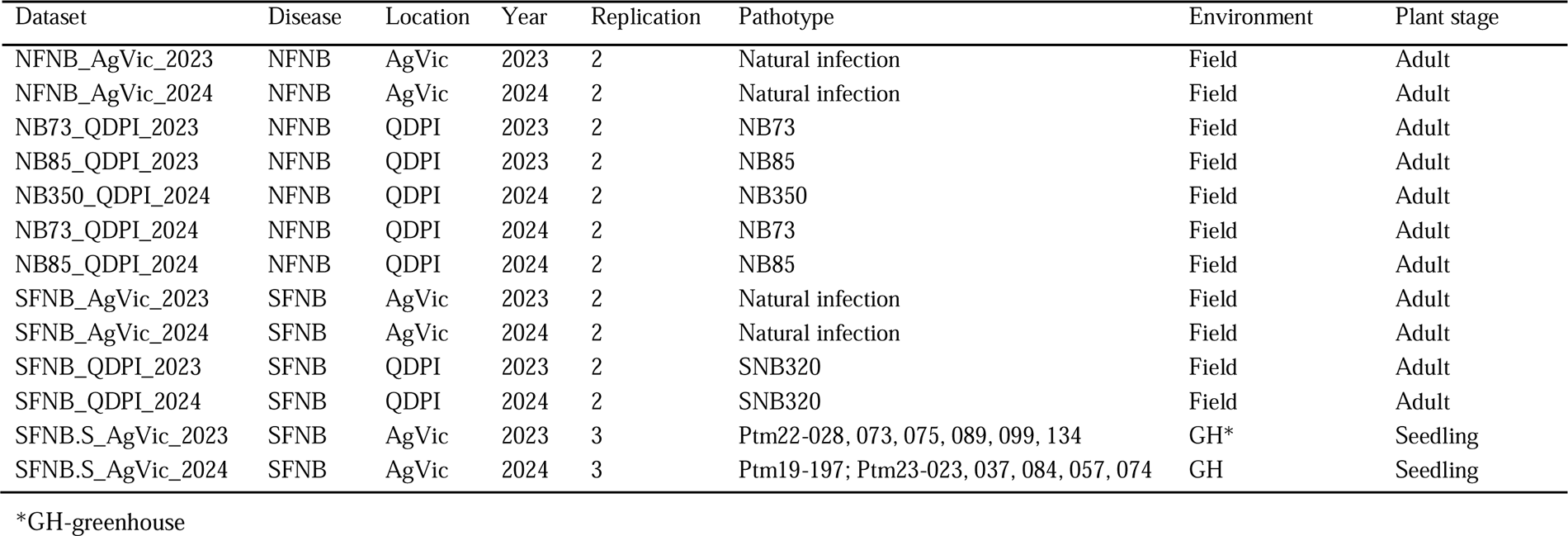
Summary of NFNB and SFNB phenotyping datasets.

### 2.3 Phenotypic data analysis

Phenotypic data for NFNB and SFNB responses were collected from 13 independent datasets capturing multiple years, locations, and disease pathotype variations (Table 1). For each experiment, a linear mixed model was fitted using ASReml-R [38] to estimate the best linear unbiased estimates (BLUEs) of genotypic performance. In the model, genotype was treated as a fixed effect, while column, row, and replicate were included as random effects. The resulting BLUEs for each experiment were used for subsequent statistical and haplotype-based mapping analyses.

Phenotypic variation and pairwise Pearson’s correlations among the 13 datasets were computed using R software [39]. Broad-sense heritability (*H²*) for each experiment was estimated as:

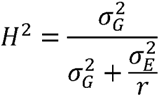

where σ^2^*_G_* is the genotypic variance, σ*^2^_E_* is the residual variance, and *r* represents the average number of replicates per genotype within each experiment.

To estimate genotype performance across experiments, a single-stage multi-environment trial analysis were performed in ASReml-R. For each NFNB and SFNB, linear mixed models were fitted to incorporate experimental design factors and relevant sources of variation. Genotype was included as a fixed effect, while location, year, pathotype, growth stage, and their interactions with genotype were treated as random effects. Several alternative model structures were evaluated by progressively adding interaction terms such as location:ID, year:ID, pathotype:ID, stage:ID, and location:year:ID to capture genotype × environment interactions. Model performance was compared using the Bayesian Information Criterion (BIC) and likelihood ratio tests (LRT), and the best-fitting model was selected accordingly. The resulting genotype BLUEs for each disease were used for downstream local genetic correlation and *in silico* stacking analyses.

### 2.4 Genotyping and marker curation

All AGG barley accessions were genotyped using the barley 40K SNP array [40], which yielded *n* = 12,084 SNP markers, where allele calls 0 = homozygous reference, 1 = heterozygous, 2 = homozygous alternate allele. Flanking sequences of all SNP markers were aligned to the Morex V3 reference genome assembly using the Blast Zone tool in TBtools [41]. Quality control and marker filtering were performed in R. Markers with >10% missing data and >10% heterozygosity were removed. After filtering, a total of *n* = 9,114 high-quality polymorphic SNP markers were retained for downstream analyses.

### 2.5 Population structure analysis

Based on the filtered high-quality markers, pairwise genetic distances among accessions were calculated using Rogers’ distance, generating a genetic dissimilarity matrix. Classical (metric) multidimensional scaling (MDS) was then applied to this matrix to perform principal coordinates analysis (PCoA). The proportion of genetic variation explained by each axis was derived from the corresponding eigenvalues. The first 23 principal coordinates, cumulatively explaining approximately 90% of the total genetic variation, were retained for downstream clustering analyses. The optimal number of clusters was determined using the NbClust package [42], which evaluates 30 clustering indices to identify the most strongly supported *K* value. Based on this result, a *k-means* clustering algorithm was applied to group the accessions into optimum *k =* 4 clusters. Three-dimensional visualization of the first three principal coordinates was generated using the R package scatterplot3d.

To visualize genome-wide relatedness patterns, the Rogers’ distance matrix was displayed as a heatmap. Rows and columns were reordered using hierarchical clustering with complete linkage, as implemented in the base R heatmap function. Cluster assignments from the *k-means* analysis were displayed as color annotations on both the rows and columns of the heatmap to illustrate the correspondence between inferred clusters and genetic dissimilarity patterns.

Pairwise linkage disequilibrium (LD) was estimated as *r*^2^ between SNP markers, and marker pairs separated by up to 10 Mb were retained for LD decay analysis. Mean *r*^2^ values were calculated within 50 kb distance bins, and LD decay curves were visualized by plotting the average *r*^2^ against physical distance.

### 2.6 Haploblock construction and localGEBV analysis

From the *n* = 9,114 high-quality SNP markers, genome-wide haploblocks were constructed based on LD using the package ‘Selection Tools’. SNP markers were grouped into LD blocks according to pairwise *r^2^* (*r^2^*≥ 0.5), with a tolerance parameter (*t* = 3) applied to accommodate potential local inconsistencies in marker order or LD estimation. This procedure was repeated until all markers had been assigned to a block, and markers not in LD with any others were retained as single-marker blocks. In total, 4,497 haploblocks were constructed across the barley genome.

To identify genomic regions with high impacts on disease response, local genomic estimated breeding value (localGEBV) analysis was performed following the framework described by Shaffer et al. (2026) [28] and Voss-Fels et al. (2019). For each dataset, marker effects were estimated simultaneously using a ridge-regression best linear unbiased prediction (rrBLUP) model. For each haploblock, the observed SNP allele combinations present in the population were defined as haplotypes [33]. The haplotype effect (localGEBV) was calculated as the sum of marker allele effects on the haplotype for all markers within the corresponding block. The variance of haplotype effects was then computed for each haploblock. As the haploblock variance is a robust alternative to the significance testing paradigm usually found in GWAS methodologies [28], haploblock were ranked according to their variance, and the ten blocks with the highest variance in each dataset were considered informative QTL. The localGEBV analytical pipeline can now be performed using a well-maintained and open-source R package HapSelect [43], which supports reproduction of the analyses reported here. Genomic positions of the identified QTL were visualized using the TBtools [41]and R package *chromPlot* [44].

### 2.7 Global and local genetic correlations

Global genetic correlations between NFNB and SFNB were estimated for each experiment using the bivariate GREML framework implemented in GCTA [45].

Local genetic correlations were calculated for each of the 4,497 haploblocks based on the haplotype effects (localGEBV) derived for each NFNB and SFNB.

For haploblock *i*, let a*_i_* = (*a_i1_*…, *a*_in_) denote haplotype effects for NFNB, *b_i_* = (*b*_i1_,…, *b*_in_) denote haplotype effects for SFNB, where n is the number of barley accessions.

Pearson’s correlation coefficient was calculated as:

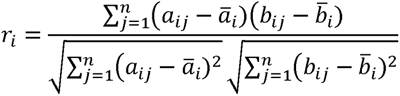

Spearman’s rank correlation was calculated as:

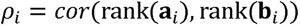

For both Pearson’s *r* and Spearman’s *p*, p-values were adjusted across all haploblocks using the Benjamini-Hochberg false discovery rate (FDR) procedure. The relationship between Pearson’s and Spearman’s coefficients was further evaluated using linear regression in the *ggplot2* package.

### 2.8 Haplotype composition analysis

To investigate patterns of haplotype composition associated with NFNB and SFNB, a joint haplotype-effect matrix was constructed by combining the haplotype effects from all identified QTL for both diseases. Hierarchical clustering was performed on the joint haplotype-effect matrix using Euclidean distance and Ward’s minimum variance method to classify accessions based on overall haplotype composition. The dendrogram was cut to group the accessions into seven clusters. Heatmaps were generated using the ComplexHeatmap package in R, with rows representing barley accessions and columns representing haploblocks. Rows were split according to the seven inferred clusters, whereas columns were separated by traits. Phenotypic disease scores for NFNB and SFNB among clusters were compared using one-way analysis of variance, followed by Tukey’s honestly significant difference (HSD) multiple-comparison test. Haplotype groups with different letters were considered significantly different at *P* < 0.05.

### 2.9 *In silico* haplotype stacking

For both NFNB and SFNB, haplotypes with negative effects were considered favourable, as they contribute to reduced disease severity. Total haplotype effects (GEBV) for each accession were obtained by summing the localGEBV across all haploblocks. Using the multi-environment BLUE values for NFNB and SFNB, a simple linear regression model was fitted to quantify the relationship between total haplotype effects and disease score.

To evaluate the cumulative contribution of favourable haplotypes, *in silico* haplotype stacking were conducted following the method described by Tong et al. (2024) [27]. The simulations were designed to estimate the potential gain in resistance by progressively replacing the original haplotypes with the favourable ones. For each disease, the ten most resistant barley accessions, as indicated by the lowest GEBV, were selected as the basis for simulation. For each selected accession, *in silico* genotypes were generated by replacing the original haplotype at each haploblock with the lowest-effect one. Haploblocks were ranked by their variance, and replacements were performed sequentially from the highest- to the lowest-variance blocks. At each step, 0.1% of all haploblocks were replaced with their most favourable haplotypes, and the total predicted haplotype effect was recalculated. The fold change in resistance was denoted by GEBV comparisons of each pair of the simulated and original genotypes [27].

To investigate the potential for improving resistance to both diseases, we further performed dual-disease stacking simulations. For each haplotype within a haploblock, its estimated effects on NFNB and SFNB were summed to form an equally weighted combined resistance score. The haplotype with the lowest summed effect was defined as the most favourable haplotype for the two diseases, and the original haplotypes were progressively replaced using the same procedure described above.

## 3 Results

### 3.1 Phenotypic variation and heritability

NFNB and SFNB were evaluated in seven and six experiments, respectively. As shown in Figure 1, a high variation in disease response was observed for both diseases. For NFNB, the field experiments in 2023 showed a distribution skewed toward resistance, likely reflecting environmental conditions favoring lower disease pressure in that season. SFNB exhibited a similar distribution pattern between field and glasshouse experiments, and overall disease severity was higher than that of NFNB.

Broad-sense heritability (*H²*) was generally high across experiments (Table S2). For NFNB, *H²* ranged from 0.60 to 0.94 (mean = 0.82), whereas SFNB showed *H²* values from 0.52 to 0.85 (mean = 0.76). The heritability of most experiments was greater than 0.75, indicating high reliability of phenotypic data. Notably, heritability was comparatively moderate in the 2023 QDPI trials, consistent with the phenotypic distribution skewed toward resistance, suggesting dynamic disease pressure and the potential influence of environmental factors (Figure 1).

**Figure 1.**
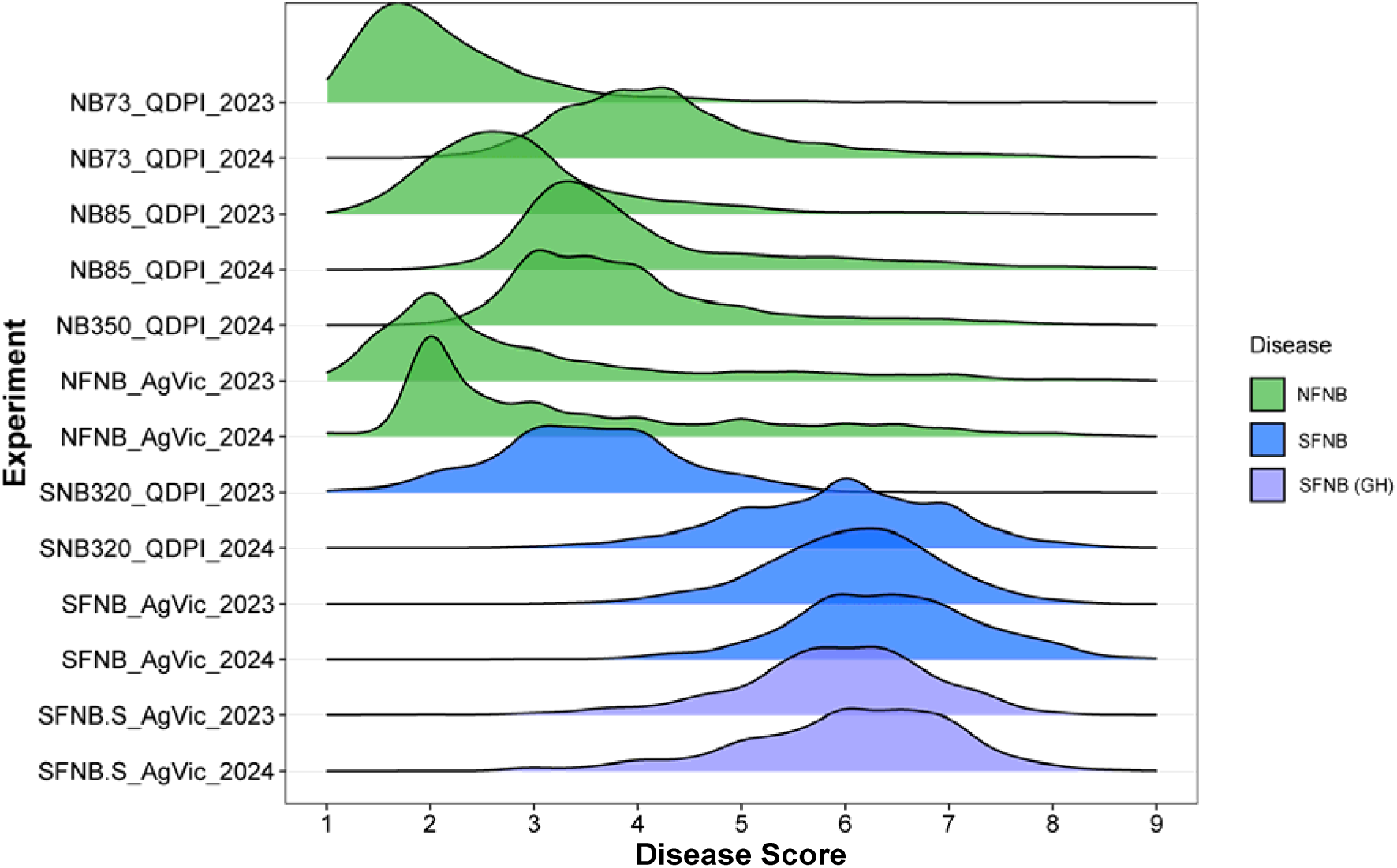
Phenotypic distribution of NFNB and SFNB across 14 experiments. Best linear unbiased estimations (BLUEs) of disease responses of each environment were used.

### 3.2 Genetic diversity and population structure

The 950 barley accessions were genotyped by the barley 40K SNP array, and after quality control, 9,114 SNP markers were used for analysis (Table S3, Supplementary Figure S1). These markers were distributed across all seven barley chromosomes, with the highest number located on chromosome 5H (1,641 SNPs) and the lowest on 1H (976 SNPs).

PCoA based on Rogers’ genetic distance revealed clear genetic differentiation among accessions. PC1 and PC2 explained 31.6% and 14.9% of the total genetic variation, respectively. *K-means* clustering separated the AGG panel into four major clusters. PC1 primarily separated Cluster 4 from the other three clusters, whereas PC2 mainly differentiated Cluster 2 (Figure 2a). The four clusters showed distinct patterns in geographical origin, growth habit, and row type (Figure 2a, Supplementary Figure S2).

Cluster 1 (*n =* 270) contained accessions originating from Asia, Europe, Africa, and North America, with a relatively large proportion of North American materials. Although spring barley was predominant, it also contained the highest number of winter barley accessions among all clusters. The proportions of two-row and six-row barley were similar. Cluster 2 (*n =* 152) consisted mainly of African accessions. This group was dominated by spring barley, with a substantial presence of six-row types (*n =* 72) alongside a moderate number of two-row types (*n =* 47). Cluster 3 (*n =* 192) showed the highest genetic and geographic diversity, including accessions from all five continents, but more than half of the entries originating from Europe. This cluster largely composed of six-row spring barley. Cluster 4 (*n =* 336) consisted predominantly of Asian accessions and was the largest cluster in the panel. Almost all accessions in this group were six-rowed spring barley, highlighting a well-defined Asian genetic background (Supplementary Figure S2).

The heatmap of Rogers’ genetic distances, reordered by complete linkage hierarchical clustering, grouped the accessions into two main clusters and further confirmed the presence of four well-defined subgroups (Figure 2b), which was consistent with the PCA patterns.

**Figure 2.**
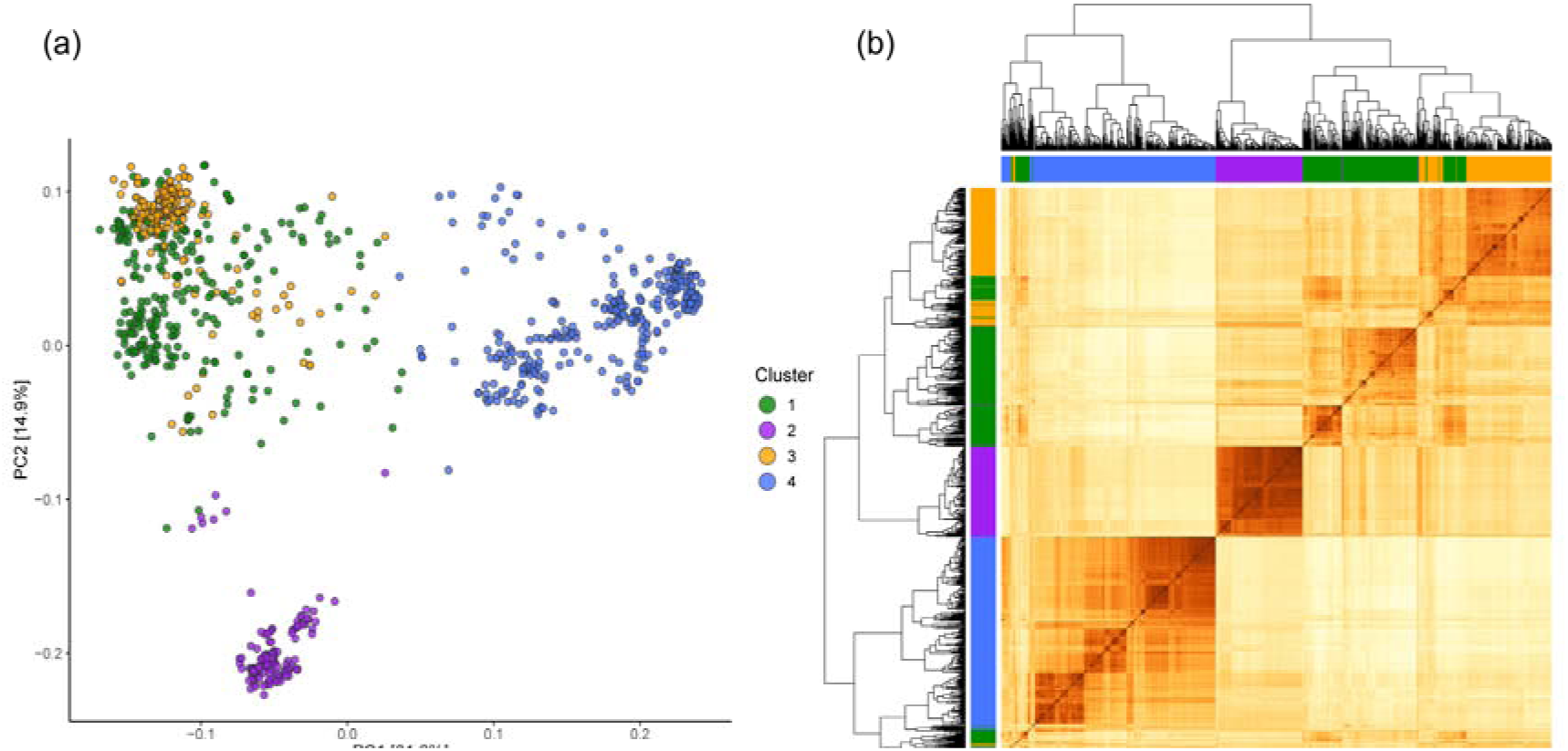
Population structure of barley accessions from AGG. (a) PCoA based on Rogers’ genetic distance matrix, with *k-means* clustering separating the AGG accessions into four clusters. The first two principal coordinates (PC1 and PC2) explain 46.5% of the total genetic variation. (b) Hierarchical clustering of the AGG population using the Rogers’ distance matrix, illustrating genome-wide relatedness among accessions.

### 3.3 Genome-wide haploblock architecture

The LD decay analysis showed that the initial mean *r*^2^ was 0.35 and rapidly decreased to 0.14 within 1 Mb (Supplementary Figure S3). Based on LD of the *n =* 9,114 curated SNP markers, a total of *n =* 4,497 haploblocks were constructed, corresponding to an average of two markers per block (Supplementary Figure S1, Table S3). The number of haploblocks varied across chromosomes, ranging from *n =* 494 blocks on 1H to *n =* 819 blocks on 5H, a pattern that closely mirrored the SNP distribution (Supplementary Figure S1). Haploblock size and marker composition showed substantial heterogeneity across the genome. For example, three haploblocks located on chromosomes 3H, 4H, and 5H each contained more than 100 SNPs (Supplementary Figure S1).

### 3.4 Correlation analyses between NFNB and SFNB

To dissect the relationship between NFNB and SFNB, five complementary correlation analyses were conducted, spanning phenotypic-, genetic-, and haploblock-level relationships (Figure 3).

Pairwise phenotypic correlations among experiments revealed moderate positive correlations for both NFNB (*r* = 0.20–0.71) and SFNB (*r* = 0.16–0.66). Correlations between the two diseases were low to moderate (*r* = –0.06 to 0.56) (Figure 3a), consistent with the phenotypic PCA, where NFNB and SFNB experiments forming distinct clusters (Supplementary Figure S4). Genetic correlations estimated from the genomic relationship matrix (GRM) were generally higher than phenotypic correlations, ranging from 0.39–0.93 for NFNB and 0.40–0.96 for SFNB (Figure 3b). Cross-disease genetic correlations (*r* = 0.07–0.82) were positive and consistently higher than the phenotypic correlations, indicating the presence of partially shared genetic determinants between NFNB and SFNB.

At the haploblock level, correlations based on haploblock variance were moderate to high among NFNB experiments (*r* = 0.43–0.90), indicating that genomic regions contributing to NFNB resistance were largely consistent across environments. In contrast, SFNB experiments exhibited comparably lower correlations (*r* = 0.23–0.67), reflecting its greater environmental sensitivity (Figure 3c). Cross-disease block-variance correlations were consistently positive (*r* = 0.17–0.74), further supporting shared genomic contributions to resistance (Figure 3).

Whole-genome haplotype-effect correlations provided finer insight into environmental responsiveness. NFNB haplotype effects were generally stable across environments (*r* = 0.35–0.89), whereas SFNB showed a wider range (*r* = -0.49–0.69). Notably, two SFNB datasets (SNB320_QDPI_2023 and SFNB.S_AgVic_2024) exhibited negative correlations with almost all other SFNB environments, despite showing high and positive block-variance correlations. Cross-disease haplotype effects correlations (*r* = - 0.49–0.78) were mostly positive, except for SFNB in SNB320_QDPI_2023, which showed negative correlations with all NFNB datasets.

Finally, local genetic correlations calculated for each of the 4,497 haploblocks (Supplementary Figure S5; Table S4) revealed highly concordant Pearson and Spearman estimates (*r* = 0.99). Approximately 60% of haploblocks showed positive correlations between NFNB and SFNB, while the remaining blocks showed negative or antagonistic relationships. Together, these correlation analyses reveal a mixture of shared and trait-specific genomic regions underlying NFNB and SFNB resistance, providing a strong foundation for subsequent haplotype-based QTL discovery and characterization.

**Figure 3.**
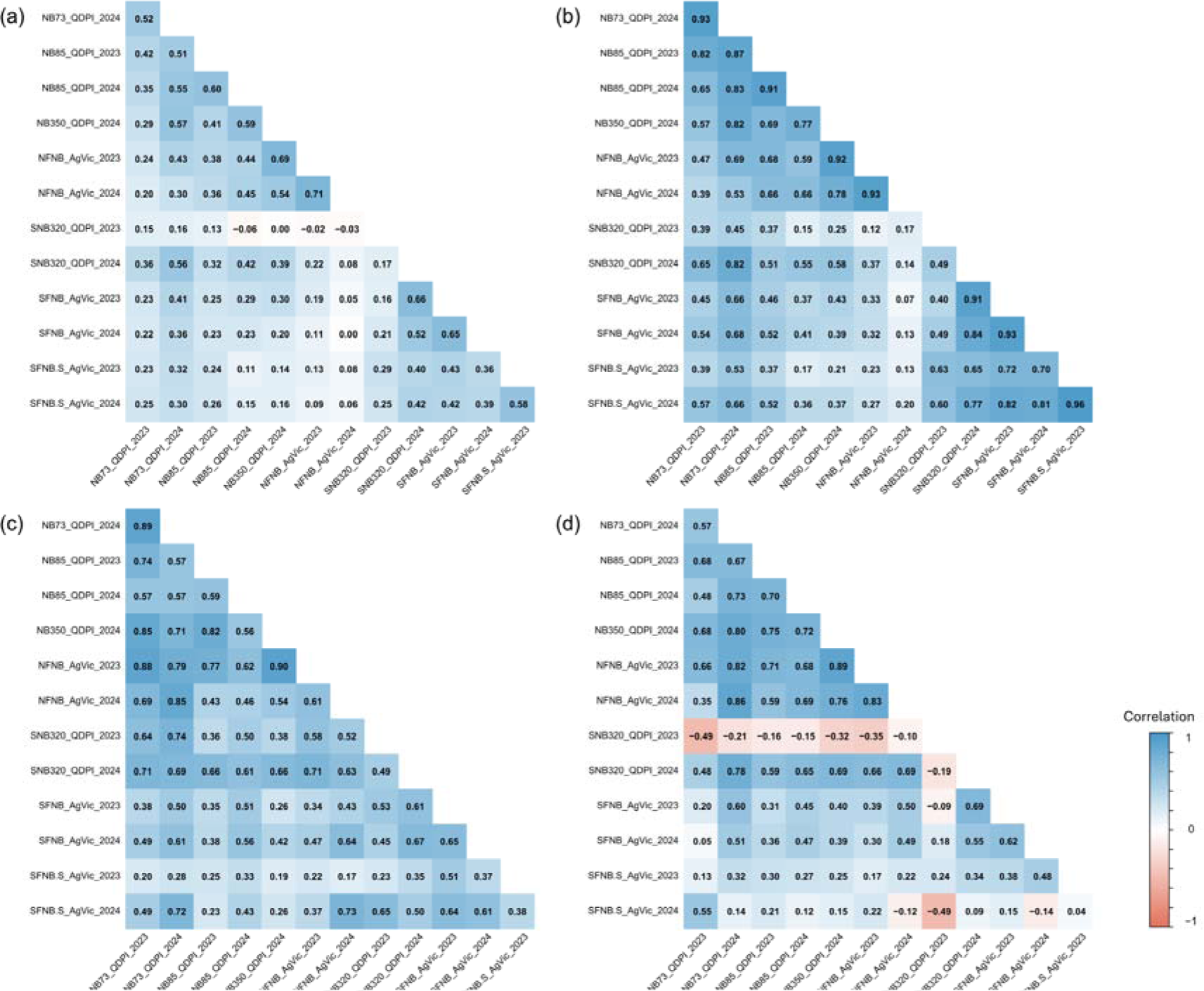
Correlations between NFNB and SFNB across individual experiments based on(a) phenotypic datasets, (b) genetic markers, (c) haploblock variance and (d) haplotype effects.

### 3.5 Haplotype-based QTL mapping

#### Overview of QTL identified across the genome

To identify genomic regions with high contribution to disease response, the ten haploblocks with the highest variance in each experiment were selected as informative QTL. This criterion has been widely used in previous haplotype-based mapping studies and is empirically supported by our data [34], where high-variance blocks consistently corresponded to large and biologically meaningful differences in haplotype effects.

As shown in Figure 4, blocks exhibiting the largest variance clearly correspond to regions with substantial divergence among haplotypes, indicating strong associations with disease response. The Manhattan plot shows that while most haploblocks have near-zero variance, a small number of blocks display distinctly elevated variance, representing putative QTL regions. Moreover, the blocks with large variances also harbour the widest spread of haplotype effects, where negative values correspond to resistance haplotypes and positive values correspond to susceptibility haplotypes (Figure 4b). Thus, the top-variance haploblocks can reliably capture major genetic signals and serve as robust candidates for QTL discovery (Figure 4, Supplementary Figure S6).

**Figure 4.**
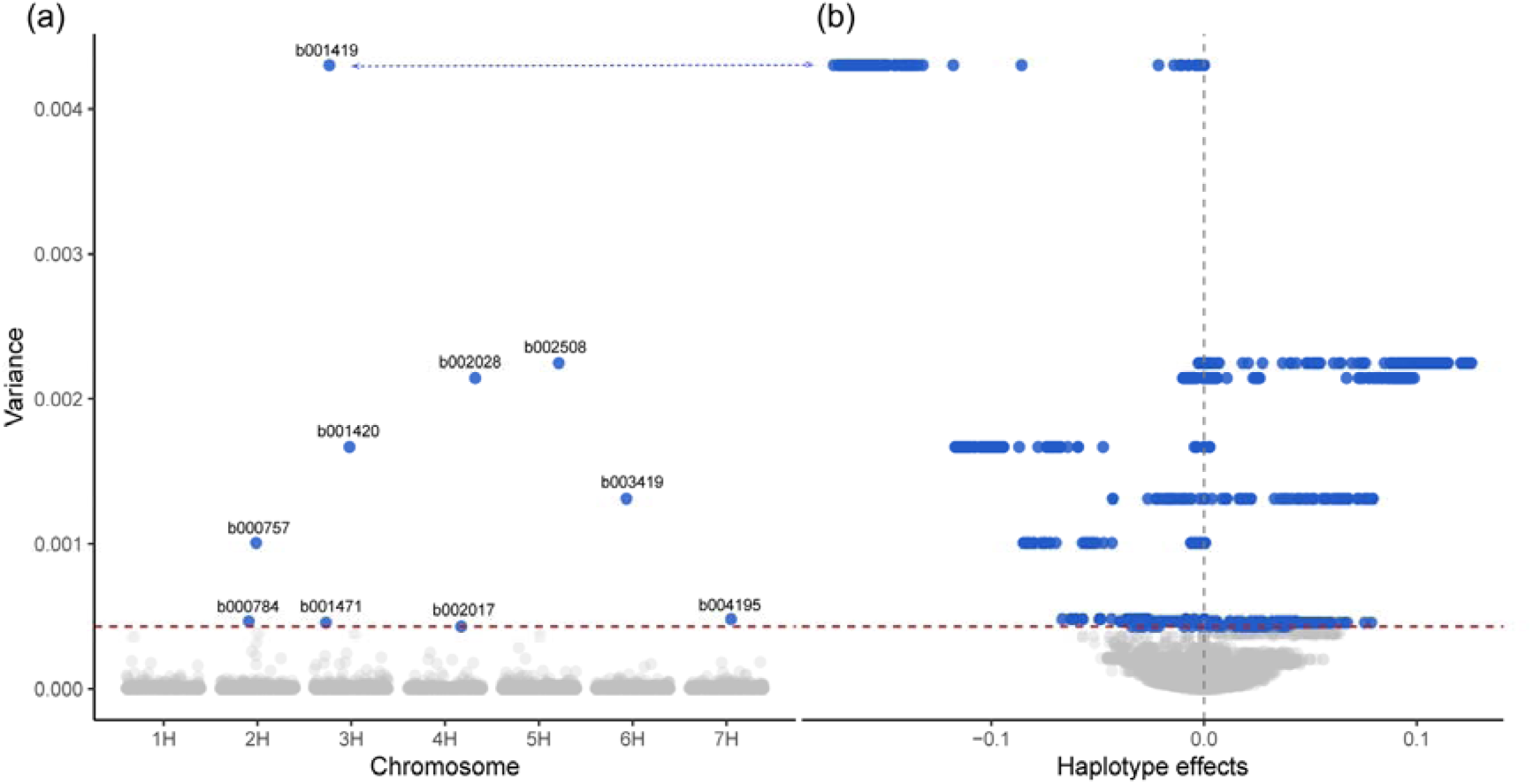
Genome-wide distribution of haploblock variance and its relationship with haplotype effects for NFNB. **(a)** Variance of haplotype effects for each of the 4,497 haploblocks across the seven barley chromosomes. **(b)** Relationship between haploblock variance and haplotype effects. Each point represents a haplotype within a block. Haplotypes with negative values were associated with reduced disease scores and therefore indicate resistance haplotypes, while positive values represent susceptibility. The top ten high variance blocks were colored blue. These results are based on haplotype-based analysis using the multi-environment BLUE values for NFNB.

For all 13 datasets, a total of 40 QTL were identified from the 4,497 haploblocks, which were associated with resistance/susceptibility to one or two diseases (Figure 5, Supplementary Figure S7, Table S5). These QTL were distributed on all seven barley chromosomes, 2H and 3H have the highest number of QTL (*n* = 9), while 6H has the lowest (*n* = 3). In total, 26 and 29 QTL were identified for NFNB and SFNB responses, respectively, of which 15 were associated with both diseases (Figure 5, Table S5). Among all QTL identified, 25 were detected in at least two experiments, providing high confidence in their effects and suggesting these loci represent relatively stable genomic regions associated with net blotch response (Table 2).

**Table 2.** Summary of high-variance haploblocks identified in at least two experiments.

| Block ID | Chr. | Flanking markers | Position (Mb) | Number of SNP | Number of experiments | Associated diseases | References |
| --- | --- | --- | --- | --- | --- | --- | --- |
| b000094 | 1H | AVRIG00323–AVRIG00356 | 234.41–264.60 | 28 | 5 | NFNB/SFNB | NFNB [46] |
| b000095 | 1H | AVRIG00358–AVRIG00391 | 265.03–287.70 | 26 | 3 | NFNB/SFNB | NFNB/SFNB [46] |
| b000412 | 1H | AVRIG01228–AVRIG01262 | 493.29–496.75 | 24 | 2 | SFNB | NFNB [47]; SFNB [48] |
| b000735 | 2H | AVRIG02057–AVRIG02089 | 147.43–163.31 | 29 | 2 | NFNB | NFNB [49]; SFNB [46] |
| b000756 | 2H | AVRIG02175–AVRIG02222 | 206.16–223.82 | 30 | 2 | NFNB/SFNB | None |
| b000757 | 2H | AVRIG02226–AVRIG02372 | 227.56–336.07 | 52 | 8 | NFNB/SFNB | NFNB [46,50]; SFNB [48] |
| b000763 | 2H | AVRIG02433–AVRIG02484 | 369.11–393.89 | 33 | 4 | SFNB | NFNB/SFNB [46] |
| b000782 | 2H | AVRIG02570–AVRIG02598 | 437.06–449.10 | 21 | 3 | SFNB | NFNB [51]; NFNB/SFNB [46] |
| b000784 | 2H | AVRIG02621–AVRIG02683 | 457.99–476.07 | 39 | 5 | NFNB/SFNB | NFNB [52]; SFNB [48]; NFNB/SFNB [46] |
| b000798 | 2H | AVRIG02734–AVRIG36934 | 488.17–501.47 | 18 | 2 | SFNB | NFNB [50] |
| b001417 | 3H | AVRIG04335–AVRIG04381 | 186.47–196.74 | 35 | 3 | SFNB | NFNB [50]; SFNB [53] |
| b001419 | 3H | AVRIG04387–AVRIG04535 | 200.85–253.65 | 112 | 10 | NFNB/SFNB | NFNB [54]; NFNB/SFNB [46] |
| b001420 | 3H | AVRIG04537–AVRIG04655 | 253.79–323.79 | 83 | 8 | NFNB/SFNB | NFNB [55]; SFNB [53]; NFNB/SFNB [46] |
| b001431 | 3H | AVRIG04689–AVRIG04757 | 337.98–355.65 | 32 | 2 | NFNB | NFNB [47]; NFNB/SFNB [46] |
| b001471 | 3H | AVRIG04920–AVRIG04979 | 406.83–423.07 | 31 | 2 | NFNB | SFNB [56]; NFNB [46] |
| b002017 | 4H | AVRIG06456–AVRIG06483 | 53.50–64.66 | 19 | 3 | NFNB | NFNB [46,55,57,58]; SFNB [48] |
| b002028 | 4H | AVRIG06543–AVRIG06978 | 95.01–399.52 | 158 | 11 | NFNB/SFNB | NFNB [50,59]; SFNB [60,61]; NFNB/SFNB [46,62] |
| b002040 | 4H | AVRIG07015–AVRIG07049 | 419.17–435.92 | 29 | 3 | NFNB/SFNB | NFNB [63]; SFNB [60]; NFNB/SFNB [46] |
| b002504 | 5H | AVRIG08176–AVRIG08211 | 69.30–78.78 | 22 | 2 | NFNB | NFNB [23,50] |
| b002508 | 5H | AVRIG08291–AVRIG08566 | 113.39–271.35 | 173 | 11 | NFNB/SFNB | NFNB [23,57]; NFNB/SFNB [46] |
| b002567 | 5H | AVRIG08782–AVRIG08808 | 374.23–380.94 | 21 | 4 | NFNB/SFNB | None |
| b003419 | 6H | AVRIG10818–AVRIG10864 | 115.65–126.44 | 35 | 8 | NFNB/SFNB | NFNB [46,64,65] |
| b004181 | 7H | AVRIG12854–AVRIG12938 | 241.84–300.57 | 65 | 4 | NFNB/SFNB | SFNB [48]; NFNB/SFNB [46] |
| b004190 | 7H | AVRIG13078–AVRIG13109 | 388.12–398.54 | 20 | 2 | NFNB/SFNB | None |
| b004195 | 7H | AVRIG13151–AVRIG13180 | 418.49–429.35 | 26 | 6 | NFNB/SFNB | NFNB/SFNB [46,62] |

Physical positions of the haploblocks were based on Morex V3 reference genome. Co-localization with previously reported QTL was assessed using the QTL catalogue compiled by Liu et al. (2026) [26].

#### Identification of novel QTL

Among the 40 QTL identified in this study, 34 overlapped with previously reported loci for NFNB or SFNB responses, whereas six were considered putatively novel (Figure 5, Table S5). These novel loci were located on chromosomes 2H (*n* = 2), 5H (*n* = 2), 6H (*n* = 1) and 7H (*n* = 1), representing previously uncharacterized genomic regions associated with net blotch resistance (Figure 5, Table S5). Blocks 2H:b000731, 5H:b002539 and 6H:b003402 were each detected in a single experiment. The remaining three potentially novel blocks, including 2H:b000756, 5H:b002567, and 7H:b004190, were associated with resistance to both NFNB and SFNB. Notably, positive local genetic correlations between the two diseases were observed at 2H:b000756, 5H:b002567, highlighting their potential value for resistance breeding, whereas a negative genetic correlation was observed at 7H:b004190 (Table S5).

**Figure 5.**
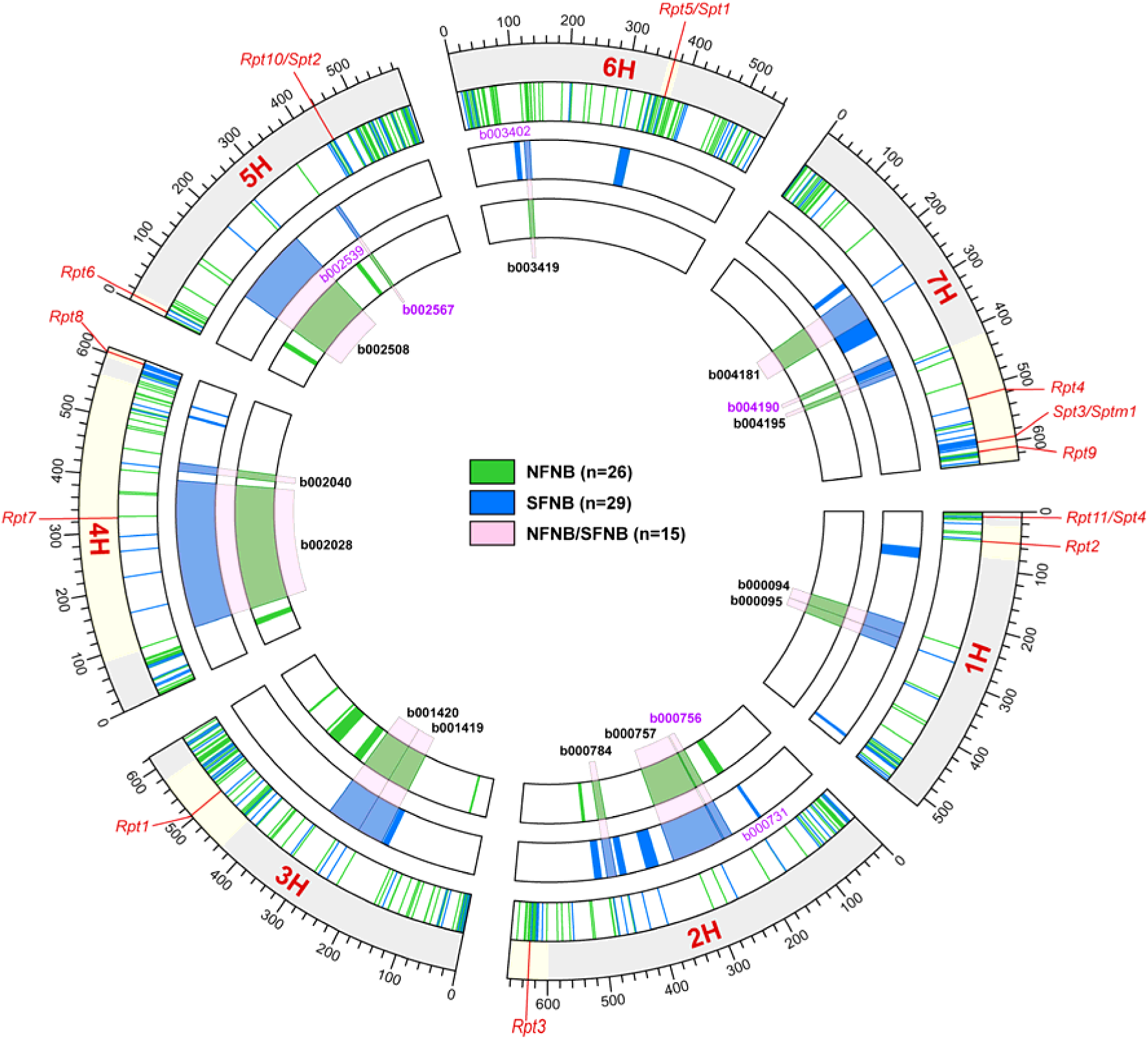
Circos plot showing 40 QTL associated with responses to net form net blotch (NFNB) and spot form net blotch (SFNB). The green segments in the inner circle represent QTL associated with NFNB, while blue segments in the second circle represent QTL associated with SFNB. Pink segments spanning both circles indicate common QTL for the two diseases, with haploblock names labeled. Potential novel QTL are highlighted in purple. The outer two circles represent previously reported QTL and major genes associated with NFNB and SFNB.

### 3.6 Haplotype composition analysis

Based on the BLUE values for NFNB and SFNB, we analyzed the haplotype composition for each QTL. Results showed that many blocks contained both resistant and susceptible haplotypes, while some only conferred resistance or susceptibility (Supplementary Figure S8). For example, block 3H:b001431 was associated with susceptibility to NFNB but resistance to SFNB, and the local genetic correlation between the two diseases at this region was strongly negative (*r* = -0.91; Table S4). In contrast, 5H:b002567 conferred resistance to both NFNB and SFNB, and the two diseases showed a strong positive local genetic correlation at this locus (*r* = 0.94; Table S4).

To further investigate haplotype composition across the population, a heatmap was generated using the haplotype effects of all 40 blocks (Figure 6a). Hierarchical clustering separated the barley accessions into different haplotype groups, providing detailed discrimination of the haplotype composition within the population. For both diseases, haploblocks located on the left side of the heatmap tended to confer susceptibility, whereas those on the right side were generally associated with resistance (Figure 6a). Two adjacent blocks on chromosome 3H, 3H:b001419 and 3H:b001420, contributed strongly to the primary separation of accessions. Groups 1 and 7 largely lacked resistant haplotypes at these loci and were also among the most susceptible groups based on phenotypic comparisons (Figure 6). Accessions carrying resistant haplotypes at these two blocks were further separated into additional groups according to their haplotype combinations at other loci. Among them, haplotype groups 4 and 6 were both resistant to NFNB, while they showed contrasting responses to SFNB, with group 4 being susceptible and group 6 showing the strongest resistance. These results were further supported by single-environment analyses, which showed consistent differences among haplotype groups across environments for both NFNB and SFNB (Supplementary Figure S9). Therefore, group 6 represents a particularly valuable source of stable dual-disease resistance.

Accessions in group 6 (*n* = 118) mainly belonged to population structure cluster 4 and were predominantly from Asia (*n* = 114), with a few from Africa (*n* = 2) and North America (*n* = 2). In contrast, group 4 accessions (*n* = 171), which were resistant to NFNB but susceptible to SFNB, mainly belonged to population structure cluster 2 and were mostly from Africa (*n* = 140), followed by Asia (*n* = 17), North America (*n* = 9), and Europe (*n* = 5) (Supplementary Figure S9; Table S6). Overall, these results demonstrate that resistance is shaped by distinct combinations of resistant and susceptible haplotypes across multiple loci. Although group 6 already carries favourable haplotype combinations for both diseases, no group or accession appeared to possess all favourable haplotypes simultaneously, suggesting that further improvement may be possible through favourable haplotype pyramiding.

**Figure 6.**
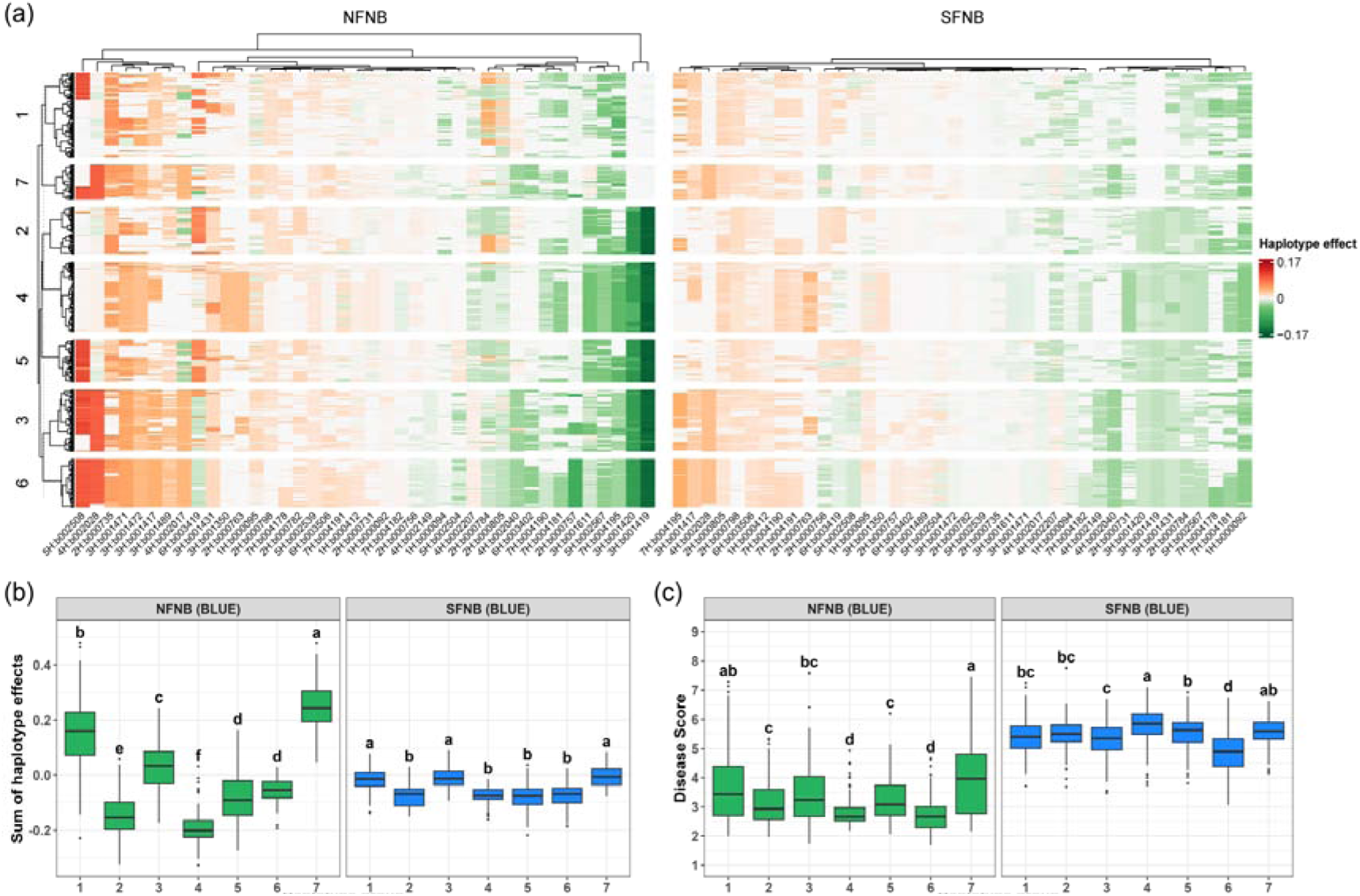
Haplotype composition across 40 QTL separated barley accessions with contrasting NFNB and SFNB resistance. **(a)** Heatmap showing haplotype effects across 40 blocks for NFNB and SFNB. Rows represent barley accessions and columns represent haploblocks. Accessions were grouped into seven haplotype groups using hierarchical clustering based on Euclidean distance and Ward’s method. Negative haplotype effects indicate resistant haplotypes, whereas positive values indicate susceptible haplotypes. **(b)** Cumulative haplotype effects across 40 QTL for NFNB and SFNB among the seven haplotype groups. **(c)** Phenotypic distributions of NFNB and SFNB disease scores among the seven haplotype groups. Different letters indicate significant differences among groups based on Tukey’s HSD test (*P* < 0.05).

### 3.7 *In silico* stacking reveals cumulative resistance potential

Across the 950 barley accessions, total haplotype effects (GEBV) showed a strong linear relationship with disease scores for both NFNB (*R*² = 0.93; Figure 7a) and SFNB (*R*² = 0.88; Figure 7b), demonstrating that the estimated marker effects and resulting localGEBV effectively capture the genetic variation in disease severity, further highlighting the robustness of haplotype-based strategies.

To explore the potential for resistance improvement, *in silico* haplotype stacking were performed. For NFNB, the ten most resistant (lowest GEBV) barley lines were selected as the basis for simulation. By replacing the original haplotypes with the most favourable ones, NFNB resistance could potentially improve by more than fourfold (Figure 7c, Table S7). A moderate improvement in SFNB resistance was also observed, consistent with the positive genetic correlation between the two diseases. For SFNB, a similar stacking strategy yielded > 4-fold improvement in SFNB resistance and an approximately 2-fold improvement in NFNB (Figure 7d, Table S8).

Next, a dual-disease stacking simulation was performed, where the favourable haplotype at each block was defined as the allele with the lowest combined effect (NFNB + SFNB). Using the ten accessions with the lowest GEBV as the basis, this simulation led to simultaneous and consistent improvements in both traits (Figure 7e, Table S9). Although the response for each disease was slightly lower than in single-trait stacking, resistance increased steadily across the stacking steps.

Importantly, dual-disease stacking was more effective for simultaneous improvement of resistance to both diseases than NFNB-only stacking. While NFNB improvement was comparable between the two scenarios (4.22-fold vs 4.54-fold), SFNB improvement was substantially greater (2.79-fold vs 1.25-fold), representing a 2.2-fold increase in SFNB gain relative to NFNB-only stacking. This illustrates that selecting haplotypes favourable to both diseases more effectively captures shared genetic components and produces dual genetic gains per replaced haploblock. In practical breeding, these results highlight the value of applying a multi-trait selection index, enabling breeders to prioritise haplotypes that simultaneously contribute to NFNB and SFNB resistance, or other important traits, thereby maximising cumulative genetic gain and facilitating the deployment of durable, broad-spectrum resistance in barley.

**Figure 7.**
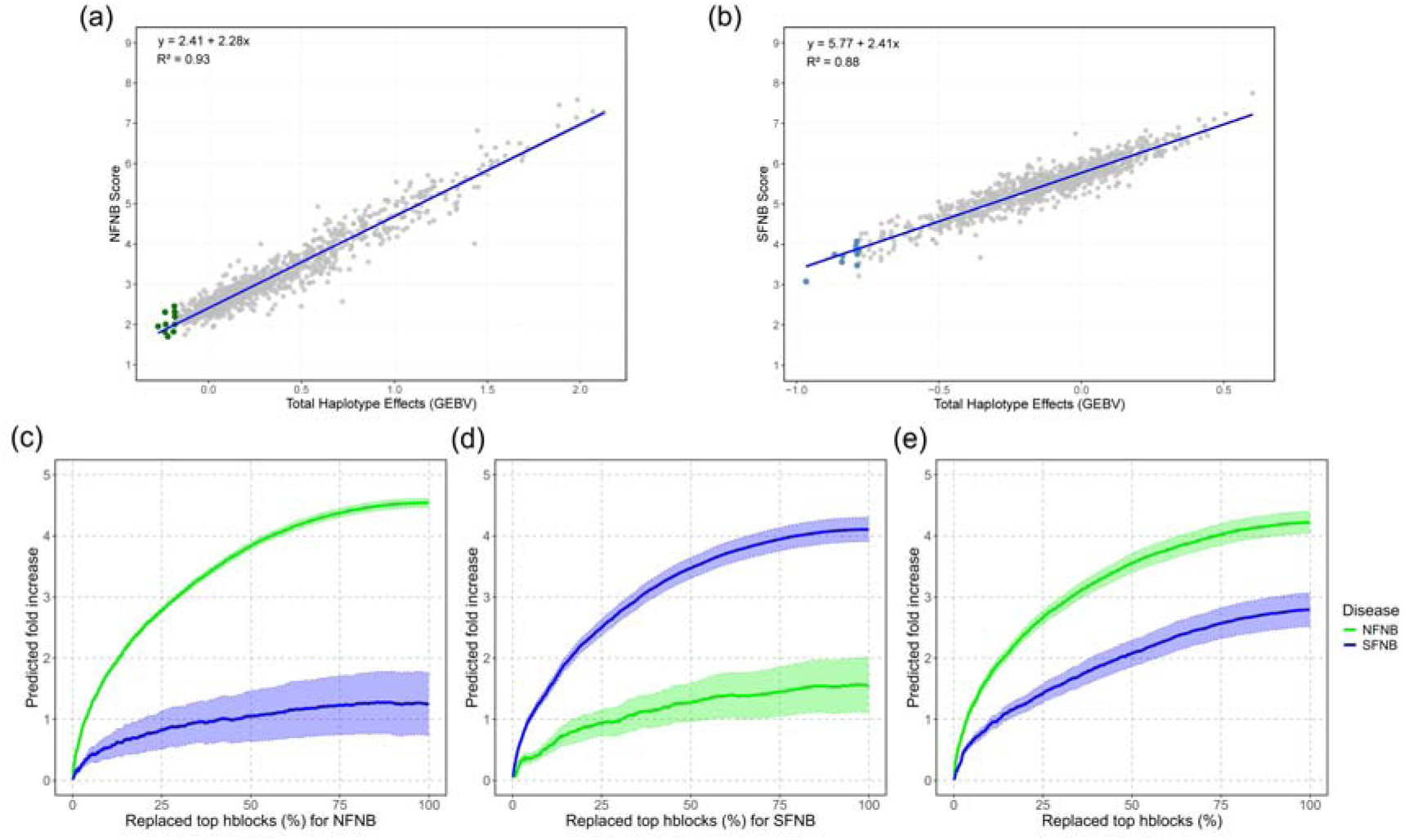
I*n silico* haplotype stacking reveals cumulative resistance potential for NFNB and SFNB. (a-b) Linear regression between total haplotype effects (GEBV) and BLUE values for NFNB (a) and SFNB (b). The colored dots indicated ten selected barley accessions used as basis of stacking simulation. (c-e) Favourable haplotype stacking simulations for NFNB (c), SFNB (d) and dual-disease resistance (e), respectively. The solid lines indicate the mean values of predicted fold changes, and shaded bands indicate standard errors.

## 4 Discussion

Using a diverse barley panel from the AGG, this study provides an integrated assessment of the genetic basis of resistance to NFNB and SFNB by haplotype-based mapping approaches. We identified important genomic regions associated with disease response, characterized the phenotypic, genetic, and haploblock-level correlations between the two diseases, evaluated population-wide haplotype composition, and assessed the potential for resistance improvement through *in silico* stacking of favourable haplotypes. These findings provide a comprehensive understanding of the genetic architecture underlying NFNB and SFNB resistance, and offer practical, haplotype-based strategies for breeding cultivars with durable and broad-spectrum disease resistance.

### 4.1 Barley AGG germplasm provided untapped genetic diversity

A major strength of this study lies in the extensive genetic diversity captured within the AGG panel. In Australia, most commercial cultivars remain susceptible to NFNB and SFNB, as consistently demonstrated by NVT disease ratings (https://nvt.grdc.com.au/nvt-disease-ratings). The narrow genetic base in elite germplasm will limit breeding progress and increase vulnerability to rapidly evolving pathogen populations. For example, in wheat, the emergence of Ug99 stem rust races has highlighted the reliance on limited resistance sources can threaten global production and food security [66]. Genebank collections therefore play a crucial role as reservoirs of allelic variation that has been lost during modern breeding. Numerous recent studies have demonstrated that plant genetic resources harbour unique resistance alleles not present in elite lines, providing opportunities to broaden the resistance base [67–69]. For barley net blotch, several GWAS conducted in diverse global panels have successfully identified novel resistance loci, further underscoring the value of leveraging genetic diversity [48,49,55–57,70–79].

The AGG panel used in this study represents one of the most diverse barley collections in Australia, compassing 950 accessions from five continents with different row types and growth habits (Table S1). This diversity resulted in substantial phenotypic variation for both NFNB and SFNB, with consistently high heritability across most experiments (Table S2), confirming the robustness and reliability of the multi-environment disease assessments. An exception was observed for the 2023 QDPI field trial, where heritability estimates were moderate (*H^2^* = 0.52–0.72). This reduction was most likely attributable to low rainfall during the growing season, which resulted in lower disease pressure across nurseries than in previous years. Such conditions are known to constrain phenotypic expression of disease resistance and reduce the power to discriminate among genotypes. This is also evident in the phenotypic PCA, where the QDPI 2023 vectors were shorter and contributed less to overall variation (Supplementary Figure S4), indicating reduced discriminatory power under low disease pressure, rather than limitations in experimental design or data quality. Nevertheless, susceptible checks remained consistently more susceptible than the other genotypes, indicating that disease development was still sufficient to preserve meaningful phenotypic differentiation.

The breadth of phenotyping further strengthened this study. For NFNB, three distinct pathotypes in addition to natural infection were evaluated, allowing the identification of broad-spectrum resistance loci effective across pathogen variation. SFNB was assessed at both the seedling and adult plant stages, providing insights into stage-specific resistance, which is essential for understanding the complex host-pathogen interactions (Table S1). Moreover, the multi-environment phenotyping of the AGG accessions provides a powerful basis for identifying environmentally stable loci and characterizing favourable haplotypes with consistent resistance effects across trials, as discussed in the following sections.

### 4.2 Phenotypic, genetic and haploblock-level relationships between NFNB and SFNB

Correlations among experiments for the same disease provide an important measure of phenotypic reproducibility and genetic stability across environments [65]. In the present study, NFNB showed consistently moderate to high correlations across experiments, particularly at the genetic and haploblock levels (Figure 3), indicating that many genomic regions contributing to NFNB resistance were stable across diverse conditions, which was consistent with previous studies [65,80,81].

In contrast, SFNB showed lower and more variable correlations across experiments, likely reflecting greater environmental sensitivity as well as increased heterogeneity in disease assessment (Figure 3). For example, the SNB320_QDPI_2023 trial, which experienced low rainfall and consequently low disease pressure, showed low but positive phenotypic correlations (*r* = 0.16–0.29), moderate genetic (*r* = 0.40–0.63) and haploblock variance correlations (*r* = 0.23–0.65) with other SFNB experiments, whereas correlations at the haplotype-effect level were weak or even negative (*r* = –0.49–0.24). Similarly, correlations between SFNB seedling and field experiments were lower and, in some cases, negative at the haplotype-effect levels. These results suggest that although similar genomic regions contributed to SFNB resistance, the direction and magnitude of haplotype effects have been strongly influenced by environmental conditions and developmental stages. Moreover, the weak correlations at the haploblock-variance and haplotype-effect levels between the two SFNB seedling experiments suggest pronounced isolate-specific genetic effects, reflecting the variability in both pathogen virulence and host resistance or susceptibility [82]. Consistent with these findings, Hassett et al. (2023) [83] analyzed the largest and most geographically diverse Australian collection of *Ptm* isolates to date using high-resolution genetic markers, revealing high genotypic diversity and low clonality within the pathogen population. Together, these results support the presence of strong isolate-dependent host-pathogen interactions, which likely contribute to the environment- and isolate-specific haplotype effects observed for SFNB resistance.

This study also provides one of the most comprehensive evaluations to date of the relationship between NFNB and SFNB (Figure 3). Previous studies have examined this relationship only relied on phenotypic correlations, which were typically weak to moderate, suggesting partial but incomplete overlap in resistance mechanisms [80,81]. By integrating phenotypic, genetic, and haploblock-level analyses, our study provides a substantially more resolved assessment of the shared and distinct genetic bases underlying resistance to the two diseases. The higher genetic correlations compared with phenotypic correlations observed in this study indicated that NFNB and SFNB share a common underlying genetic foundation, while environmental conditions and pathotype variation obscure this relationship at the phenotypic level. Consistently positive correlations at the haploblock-variance level further support the presence of genomic regions contributing to resistance against both diseases. Approximately 60% of haploblocks showed positive local genetic correlations between NFNB and SFNB, providing strong evidence for shared resistance loci (Supplementary Figure S5). However, a substantial proportion of haploblocks exhibited weak or negative correlations, indicating disease-specific effects or antagonistic loci where haplotypes conferring resistance to one disease may increase susceptibility to the other. These results are biologically plausible given that NFNB and SFNB are caused by two closely related but genetically differentiated *formae speciales* of *Pyrenophora teres* [6,7]. Despite their shared evolutionary origin, comparative genomic studies have revealed extensive structural variation, transposable element expansion, and divergence in effector repertoires between *Ptt* and *Ptm*, providing a genomic basis for the incomplete overlap in host resistance observed here [84]. These differences reflect distinct infection strategies and selection pressures acting on the two pathogen lineages, resulting in differentiation of virulence-associated loci.

### 4.3 QTL discovery by haplotype-based approach

Using the haplotype-based mapping method, we identified a total of 40 QTL associated with resistance to NFNB and SFNB in the AGG barley diversity panel, including 15 loci associated with both diseases (Figure 5, Table S5). Most of these loci co-localized with previously reported QTL, demonstrating the robustness of haplotype-based mapping and its consistency with earlier linkage mapping and GWAS studies (Table 2, Table S5). Notably, approximately half of the detected QTL overlapped with those reported by Jambuthenne et al. (2026), despite the two studies analysing different populations from AGG. Given that both studies applied the same haplotype-based methodology, the concordance highlights the reproducibility and transferability of the localGEBV approach across populations and disease datasets.

Beyond validating known resistance regions, we also identified six potentially novel QTL, underscoring both the analytical power of haplotype-based mapping approach and the untapped resistance diversity preserved within the AGG collection. For example, the block 5H:b002567 were associated with resistance to NFNB and SFNB at both seedling and adult plant stages, and most haplotypes within this block conferred resistance, indicating that it represents a favourable genomic region that can be readily exploited in breeding programs (Table S5).

Compared with single-marker associations, haplotype-defined QTL represent inherited chromosome segments that capture multi-allelic effects within LD blocks and more closely reflect functional genetic units [28]. This feature makes haplotype-based QTL particularly relevant for breeding, as these genomic segments are less likely to be disrupted by recombination and can be more reliably tracked and introgressed. Consistent with this, 25 QTL were repeatedly detected in at least two experiments, including 15 that were associated with both diseases (Table 2; Table S5). Notably, 23 of these stable QTL overlapped with loci reported in previous studies (Table 2; Table S5), providing further validation of their function in disease responses and supporting their value as high-confidence targets for resistance breeding.

In some cases, haploblocks spanned large genomic intervals, particularly in pericentromeric regions where recombination is reduced. For example, two adjacent blocks on chromosome 3H, 3H:b001419 and 3H:b001420, corresponding to a large region (200.85–323.79 Mb), were repeatedly detected in more than eight experiments and were associated with resistance to both NFNB and SFNB. The same genomic region was also detected in another AGG population, where it was associated with resistance to NFNB and SFNB [46]. While several previous studies have reported QTL in this region [53–55], most were mapped to broad regions, likely due to limited marker density and suppressed recombination rate around the centromere. In the present study and in Jambuthenne et al. (2026), this region showed major effects on resistance to multiple pathotypes, and accessions carrying the resistant haplotypes at these blocks exhibited lower disease scores (Figure 6), highlighting the value in barley resistance breeding.

In contrast, the largest blocks 4H:b002028 identified in this study, which spans more than 300 Mb on 4H (95.01–399.52 Mb), was associated with susceptibility to the two diseases. This block co-located with resistance gene *Rpt7*, as well as several previously reported QTL for NFNB, including *QNFNBAPR.Al/S-4Ha* [50], *QRpts4* [85] and *Rpt-4H-5-7* [63], and for SFNB, including *QTL6* [61] and *QRptm-4H-43-57* [60]. Notably, many of these loci were defined by broad confidence intervals exceeding 150 Mb, consistent with the reduced mapping resolution expected in low-recombination regions. Interestingly, the stem rust-related gene *Rpr1*, which was mapped to 145.53–411.43 Mb, also overlaps with this broad genomic region [86]. *Rpr1* is required for *Rpg1*-mediated stem rust resistance but is not required for resistance mediated by genes *rpg4* or *Rpg5*, indicating pathway specificity in rust resistance [86]. Another example is the block 5H:b002508, which was mapped to 113.39–271.35 Mb, conferring susceptibility to net blotch. It overlapped with the leaf rust resistance gene *Rph2*, that is currently delimited to a large 278-Mb interval (34.07–312.74 Mb) [87]. Although the biological relevance of *Rpr1* and *Rph2* to net blotch response remains unclear, their co-location with susceptibility-associated haploblocks is noteworthy. Given the distinct infection strategies of rust pathogens, which are generally biotrophic, and net blotch pathogens, which are necrotrophic, resistance mechanisms effective against one pathogen lifestyle may not necessarily confer resistance to another and may, in some cases, have contrasting effects [88]. Therefore, these broad intervals may harbor genes involved in pathogen-specific defense responses, susceptibility-associated pathways, or closely linked loci with opposing effects on different diseases. Future studies using higher-density markers, or secondary mapping populations will be necessary to resolve these broad intervals and distinguish individual causal genes from linked haplotypic effects.

### 4.4 Implications of haplotype grouping and stacking

The haplotype composition analysis demonstrated that resistance to NFNB and SFNB is determined by distinct combinations of favourable and unfavourable haplotypes across multiple QTL. Although several major loci, such as 3H:b001419 and 3H:b001420, contributed strongly to the separation of haplotype groups, the contrasting disease responses among groups indicate that resistance cannot by fully explained by single loci alone (Figure 6). Instead, resistance differences among accessions were better explained by the combined effects of haplotypes across multiple QTL. The contrasting performance of groups 4 and 6 is particularly informative. Both groups showed strong resistance to NFNB, but they differed markedly in their response to SFNB, with group 4 being susceptible and group 6 showing stable resistance across environments (Figure 6, Supplementary Figure S9). This suggests that NFNB and SFNB resistance share some genetic components but also require disease-specific favourable haplotype combinations, which was consistence with the correlations between them. Group 6 therefore represents a valuable source of stable multi-disease resistance, whereas group 4 may provide useful NFNB-specific resistance alleles that could be combined with SFNB resistance sources in future breeding (Table S6). Importantly, no haplotype group appeared to carry all favourable haplotypes simultaneously, suggesting that additional resistance potential remains untapped in the current germplasm.

The *in silico* haplotype stacking conducted in this study illustrate the cumulative genetic potential achieved by progressively replacing unfavourable haplotypes with beneficial ones (Figure 7). Although stacking thousands of haplotypes is not realistic in practical breeding, these simulations define the potential of genetic improvement and provide insights into how favourable haplotypes may be combined to maximize resistance gains. For both NFNB and SFNB, stacking resulted in a progressive increase in predicted resistance, consistent with the highly polygenic nature of net blotch resistance. A key insight from these simulations is that dual-disease stacking was more efficient than single-disease stacking, demonstrating that prioritizing haplotypes with pleiotropic effects yields greater multivariate genetic gain than focusing exclusively on single disease.

Although real-world breeding is constrained by recombination, population size, and selection intensity, the stacking framework provides a quantitative basis for prioritising which haplotypes and genomic regions are most valuable to target. With the advancement in genomic technologies, haplotype stacking insights can be readily integrated with genomic selection, selection indices, and AI-assisted optimization frameworks, such as genetic algorithms and Bayesian optimization approaches for parental and crossing scheme design [89–91]. By focusing on pleiotropic haploblocks and stacking favourable haplotypes, breeders can more efficiently assemble favourable allele combinations and accelerate progress toward barley cultivars with durable, broad-spectrum resistance to net blotch.

## 5 Conclusion

This study demonstrates that haplotype-based mapping combined with multi-environment phenotyping provides a powerful framework for dissecting the complex genetic architecture of resistance to net blotch in barley. Using a diverse panel of AGG accessions, we identified 40 QTL associated with resistance/susceptibility to NFNB and SFNB, revealing both shared and disease-specific genetic components underlying responses to the two net blotch forms. Correlation analyses further revealed that NFNB and SFNB resistance are partially governed by common genetic components, while also involving distinct disease-specific mechanisms. Haplotype composition analysis showed that resistance patterns were better explained by the combined effects of haplotypes across multiple QTL than by individual loci alone. *In silico* haplotype stacking further highlighted the potential for improving resistance to single and multiple diseases through the pyramiding of favourable haplotypes. Collectively, these findings provide new insights into the genetic basis of net blotch resistance and support the use of haplotype-informed breeding strategies to develop barley cultivars with durable and broad-spectrum resistance.

## Author’s contribution

DL analyzed the data and wrote the manuscript; ED conceptualized the study; XCZ, LS, TG, HW, HD, MM guided the design and establishment of field screening nurseries; JYT and CSC contributed to data analyses; DJG, SP, LH, BH, ED supervised the research project and critically reviewed and revised the manuscript. All authors read and revised the manuscript.

## Acknowledgments

This research was supported by the GRDC Net Blotch Consortium (UOQ2005-012RTX, UOQ2405-015RTX). DL was supported by the China Scholarship Council (202203250003) and the GRDC Research Scholarship (UOQ2601-009RSX).

## Ethical standards

This article does not contain any studies with human participants or animals performed by any of the authors.

## Data availability

The datasets generated or analyzed during the current study (i.e. genotypes, and mapping data) are available from the corresponding author on reasonable request.

## Supplementary Figures

**Figure S1.**
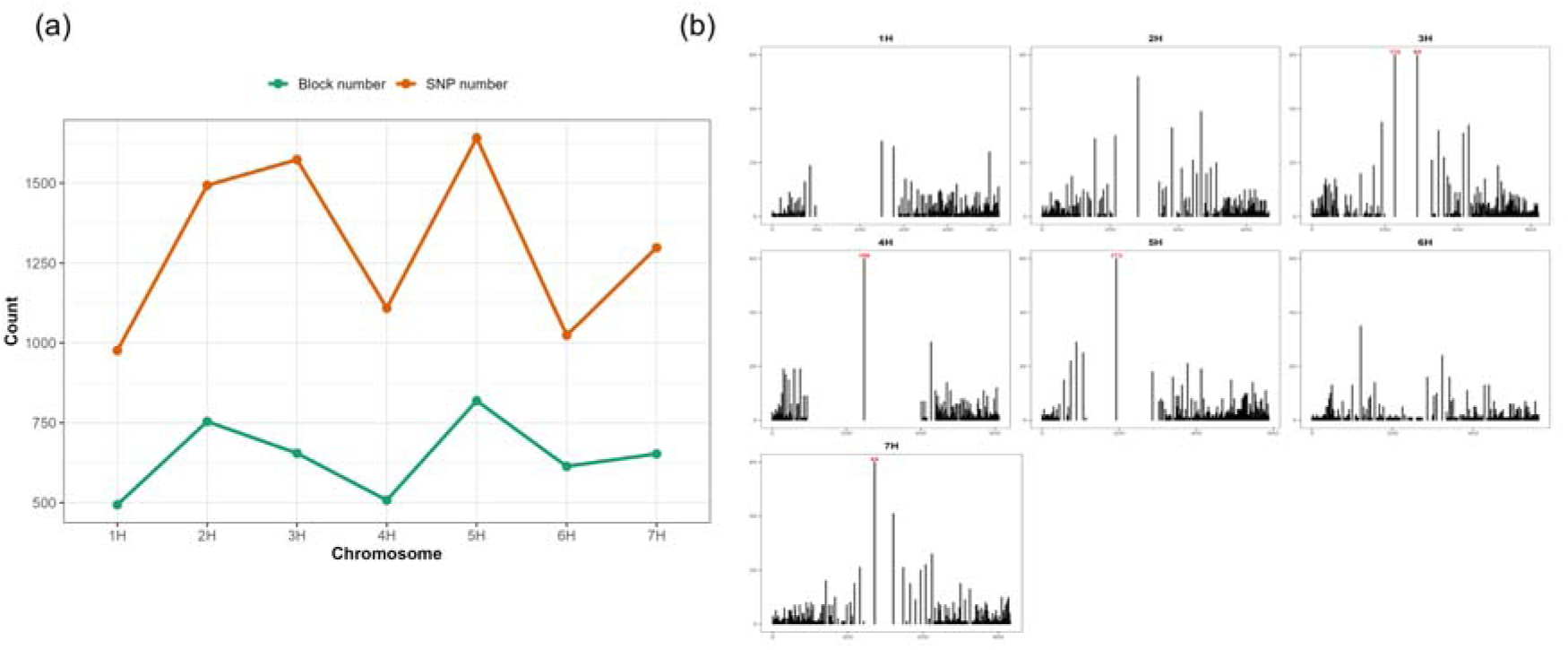
Distribution of SNPs and blocks across the barley genome. **(a)** Number of SNPs and blocks per chromosome (1H–7H). **(b)** Genome-wide distribution and size of blocks. The bars on 3H, 4H and 5H exceed the y-axis scale and represent large LD blocks comprising 112, 83, 158, 173 SNPs, respectively.

**Figure S2.**
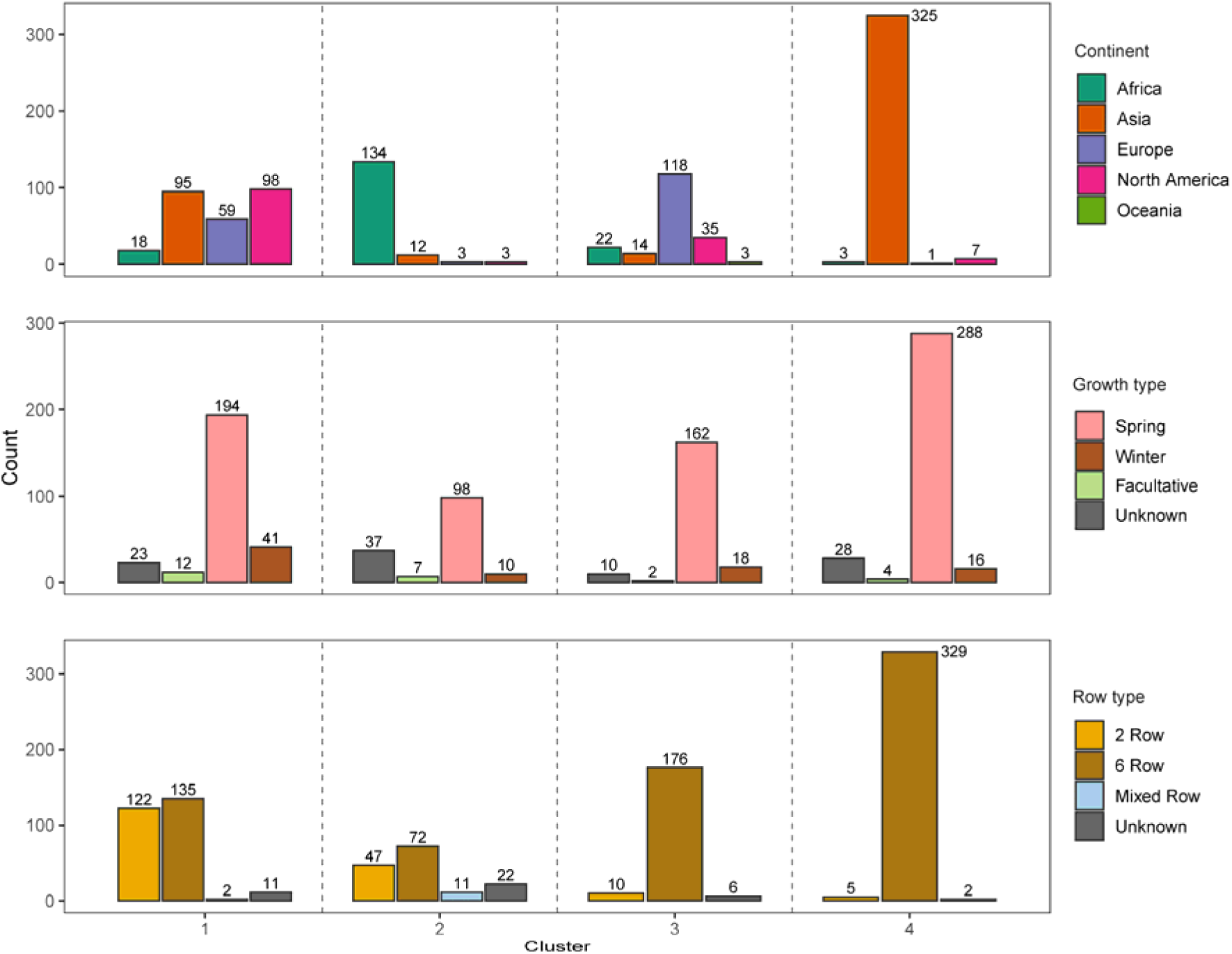
Distribution of barley accessions across continents, growth and row types in the AGG panel. Counts are displayed for each k-means cluster, with numbers above bars indicating the number of accessions.

**Figure S3.**
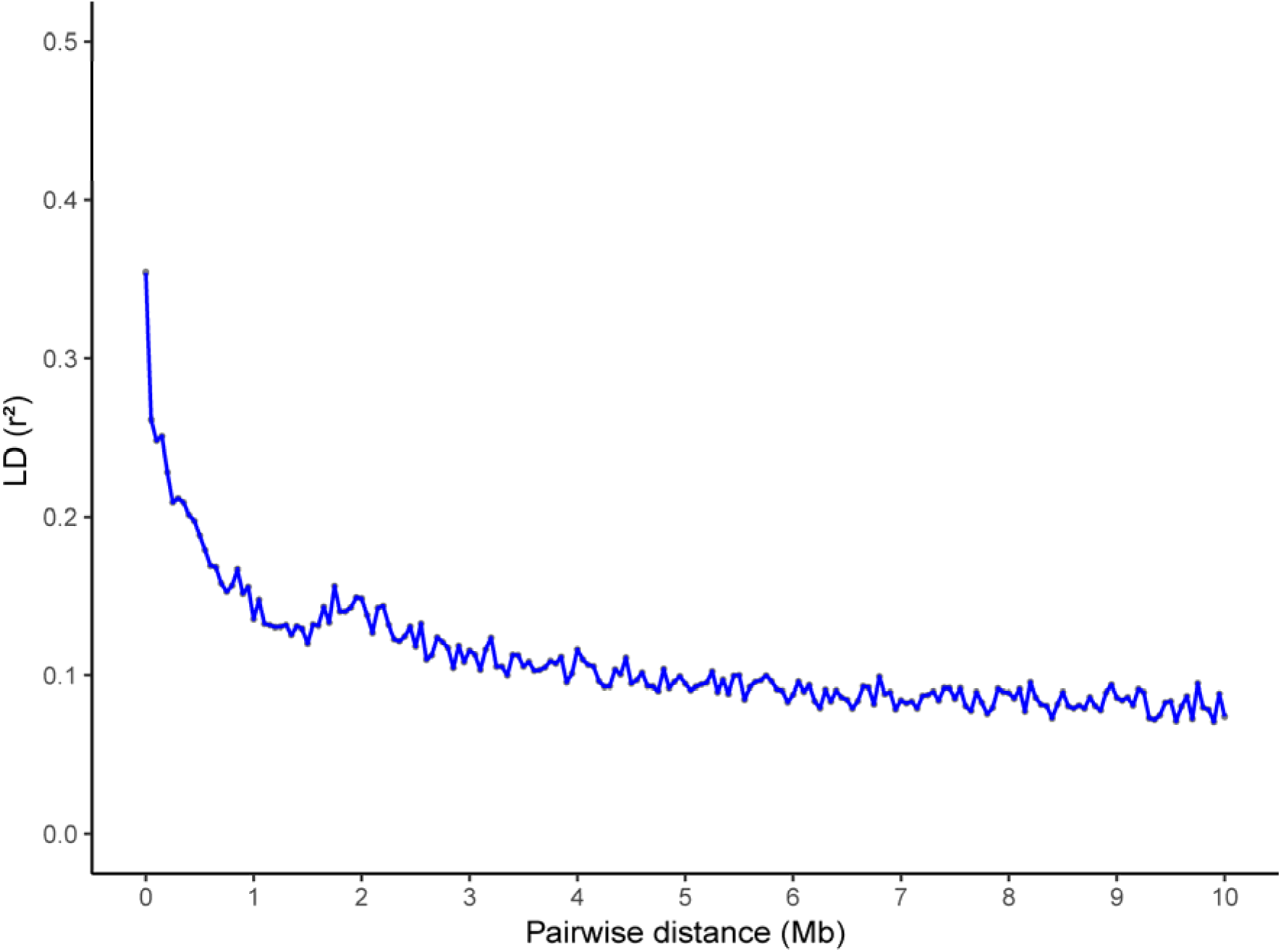
Linkage disequilibrium (LD) decay over physical distances (Mb). Pairwise LD was estimated as *r*^2^ between SNP markers, and mean *r*^2^ values were calculated within 50 kb distance bins.

**Figure S4.**
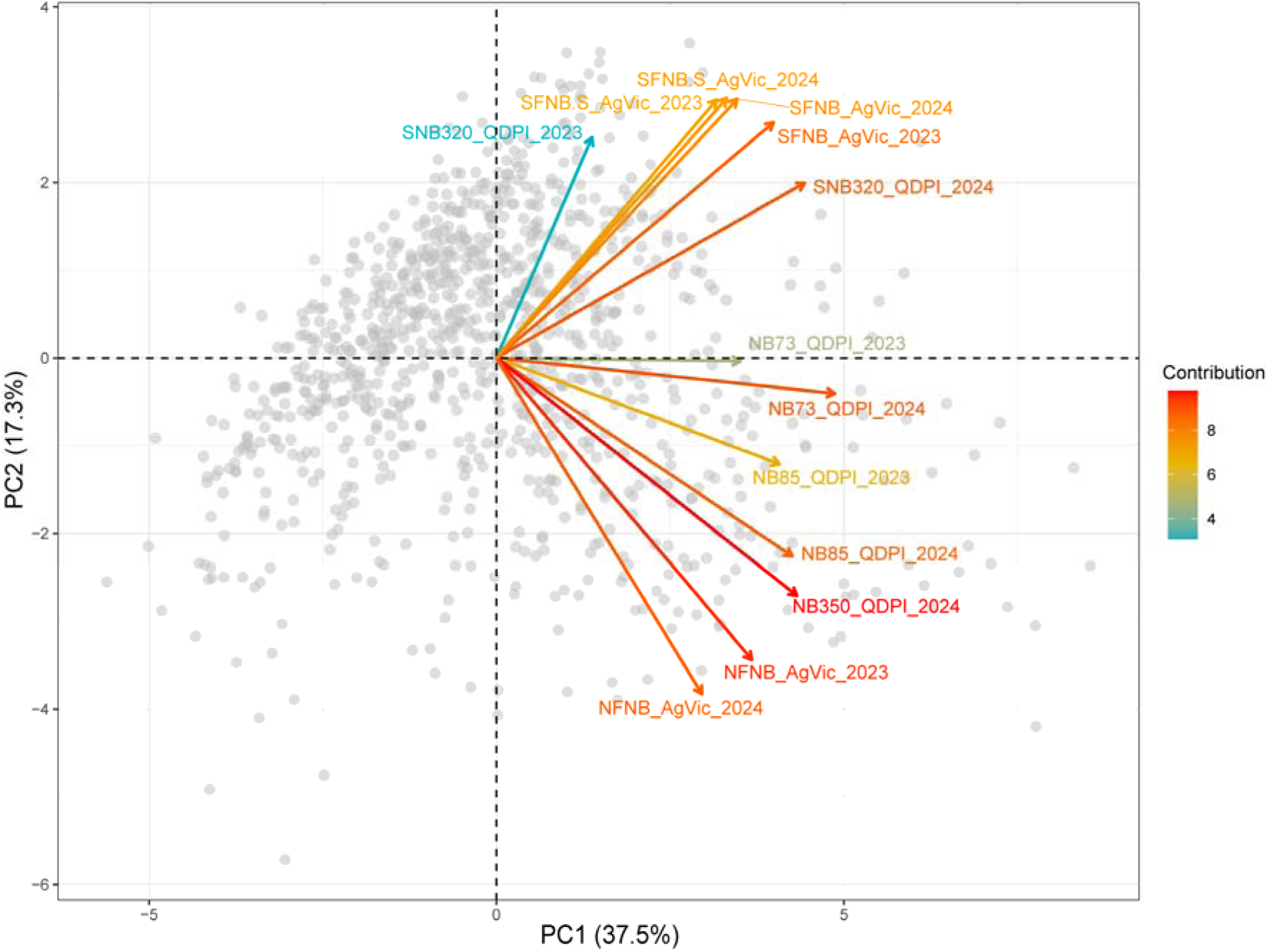
Phenotypic principal component analysis (PCA) of NFNB and SFNB across 13 experiments. Colored arrows indicate environmental variables, with arrow length reflecting the strength of their contribution to the principal components; gray dots are the entire AGG accessions; the scale on the right represents the variable’s contributions to the component.

**Figure S5.**
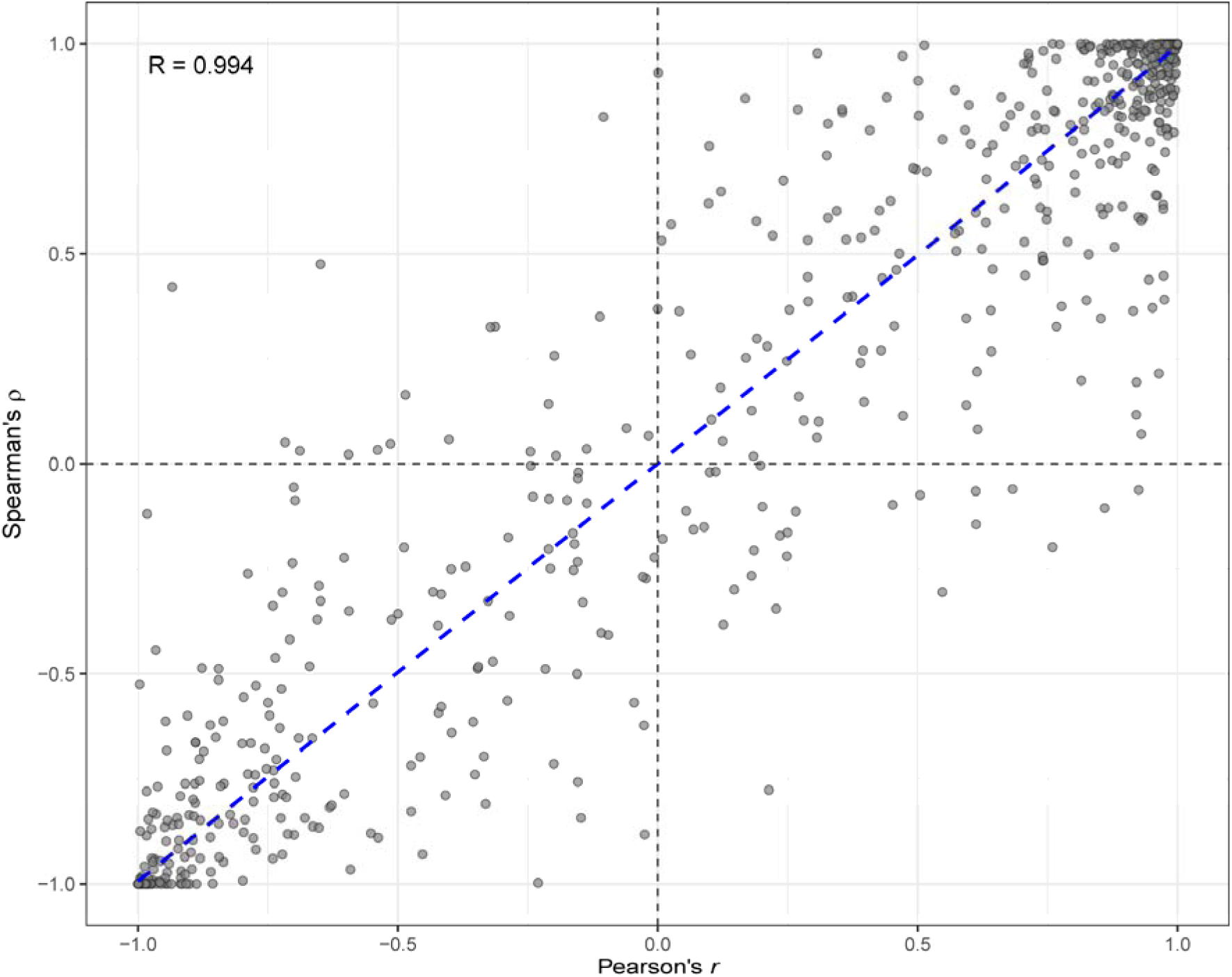
Local genetic correlations at the haploblock level. Each dot represents a single haploblock.

**Figure S6.**
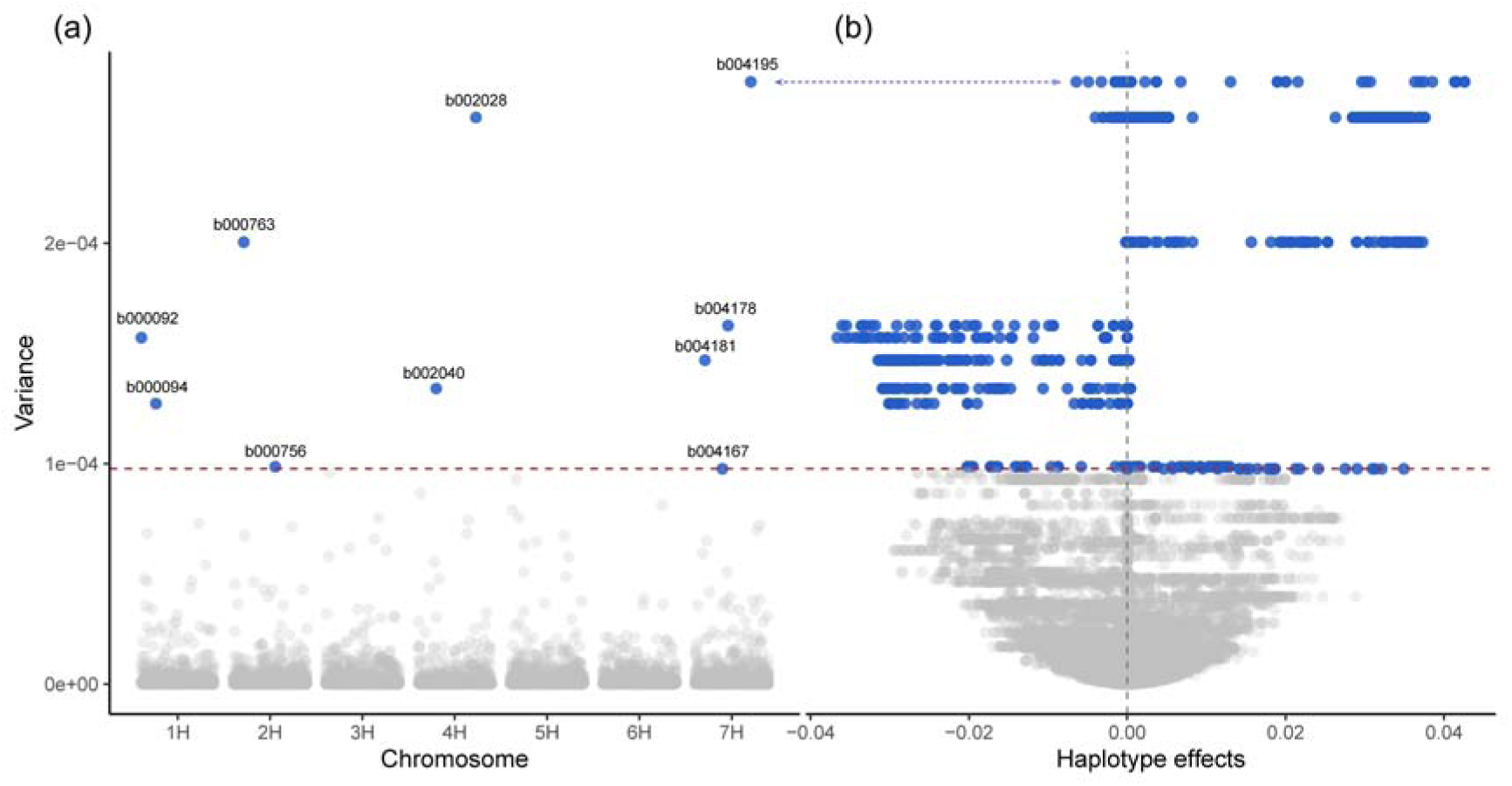
Genome-wide distribution of haploblock variance and relationship with haplotype effects for SFNB. **(a)** Variance of haplotype effects for each of the 4,497 haploblocks across the seven barley chromosomes. **(b)** Relationship between haploblock variance and haplotype effects. Each point represents a haplotype within a block. Haplotypes with negative values were associated with reduced disease scores and therefore indicate resistance haplotypes, while positive values represent susceptibility. The top ten high variance blocks were colored blue. These results are based on haplotype-based analysis using the multi-environment BLUE values for SFNB.

**Figure S7.**
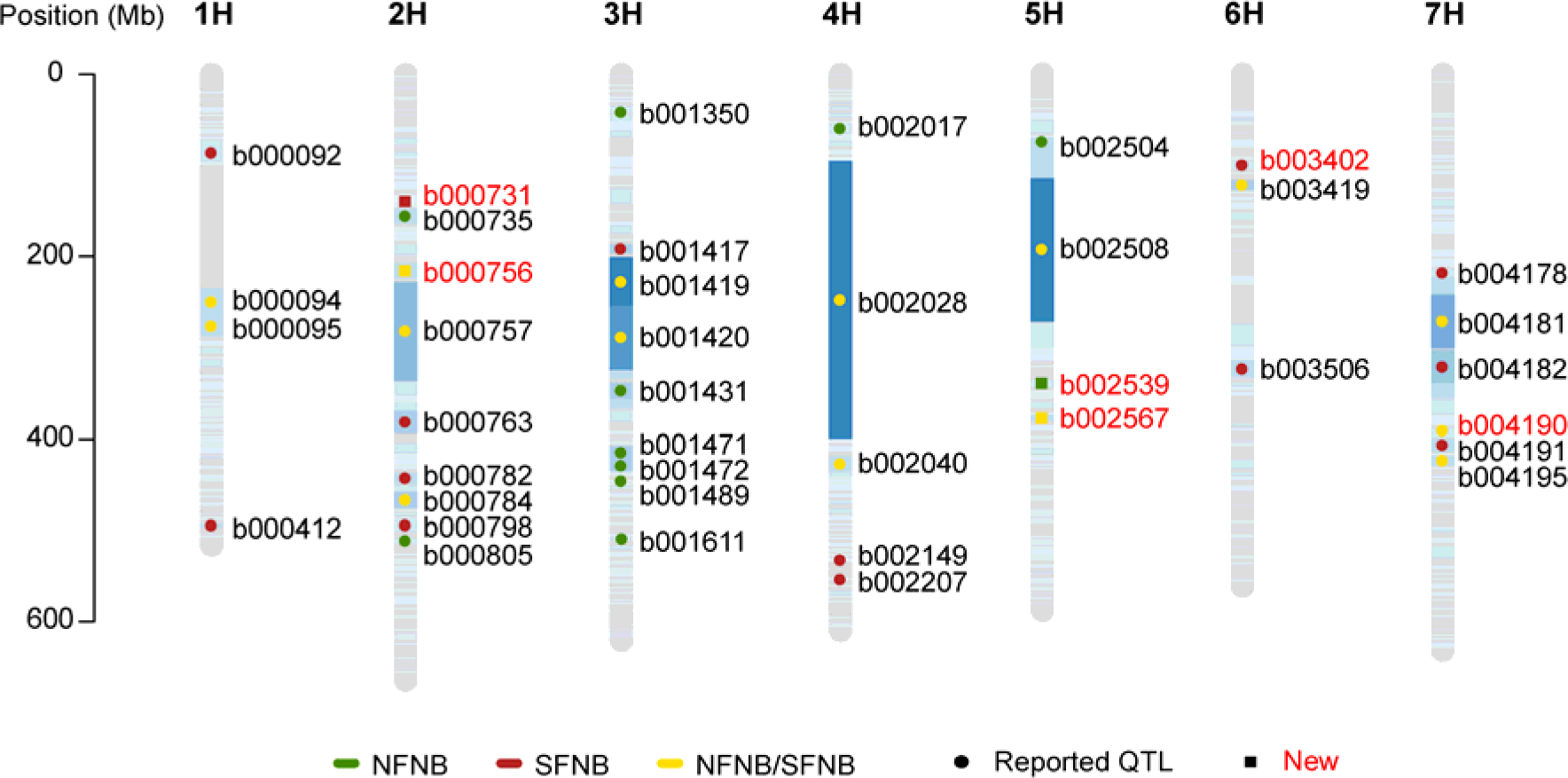
Genome-wide distribution of NFNB and SFNB QTL identified in 13 datasets. In each chromosome, blue gradient bars represent haploblocks, with darker shades indicating greater SNP number. Chromosomal positions (Mb) are shown on the left. Colored markers denote QTL associated with resistance to net form net blotch (NFNB; green), spot form net blotch (SFNB; dark red), and loci detected for both diseases (gold). Previously reported loci were shown as circles, whereas new QTL identified in this study are shown as squares and highlighted with red labels.

**Figure S8.**
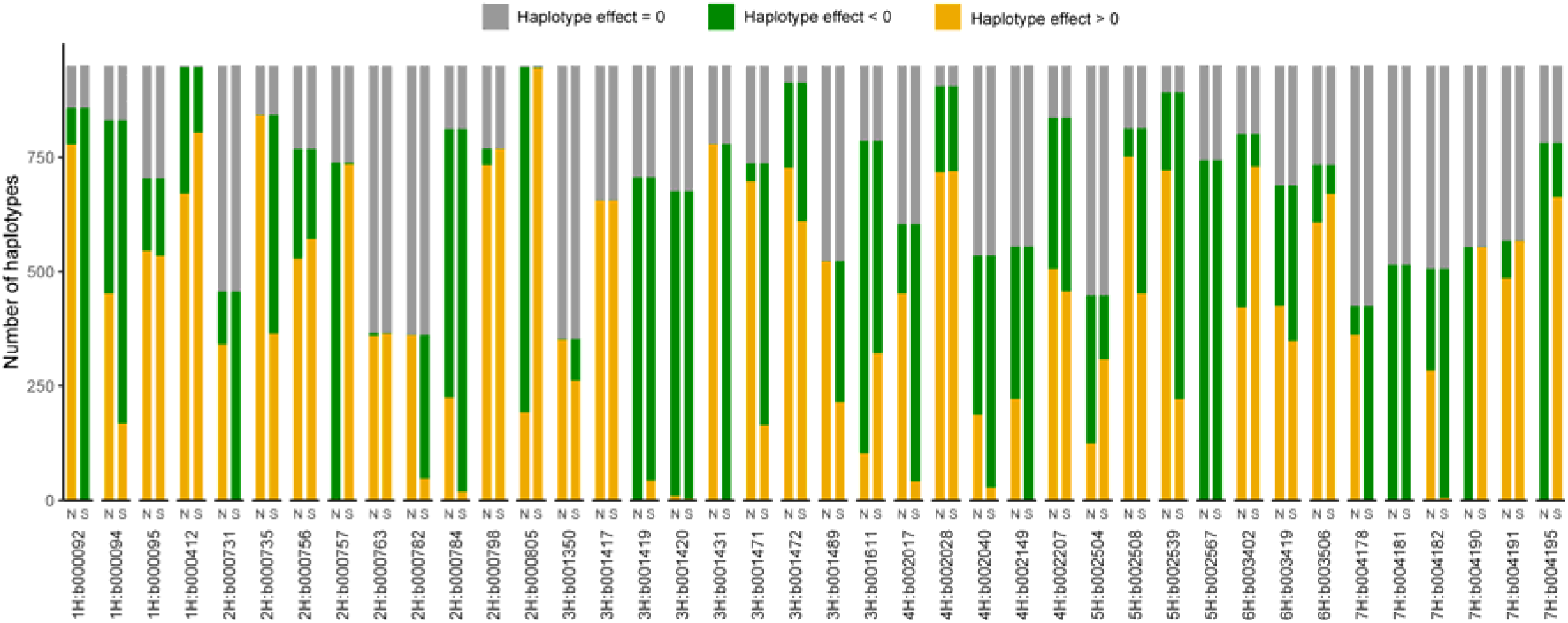
Distribution of haplotype effect classes across 40 QTL for NFNB (N) and SFNB (S). Stacked bar plots show the number of accessions carrying resistant (Haplotype effect < 0), neutral (Haplotype effect = 0), and susceptible (Haplotype effect > 0) haplotypes within each block.

**Figure S9.**
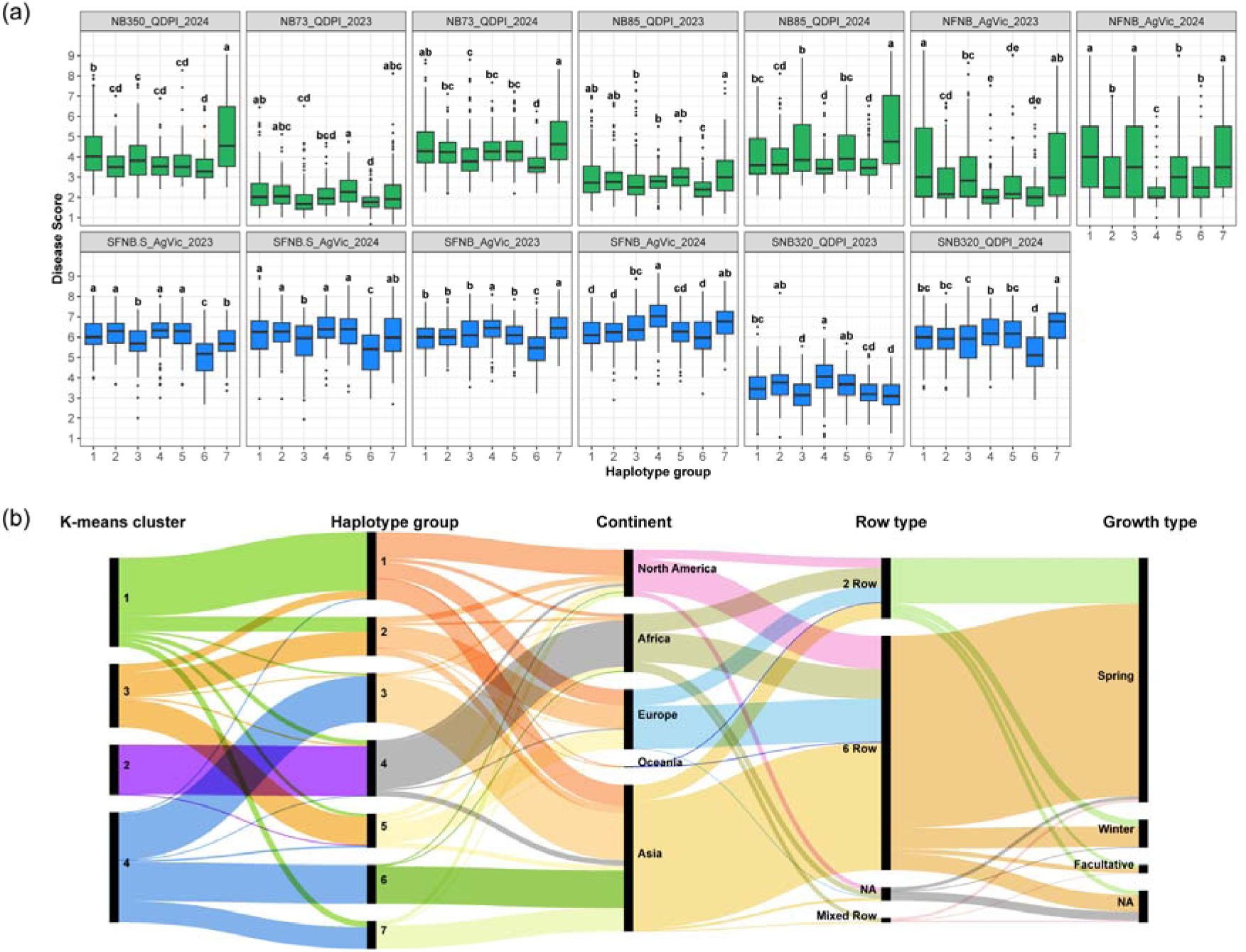
Phenotypic effects and passport information of the seven haplotype groups. **(a)** Phenotypic differences among the seven haplotype groups across 13 experiments. Boxplots show disease score distributions for each haplotype group within each experiment. Different letters indicate significant differences among groups based on Tukey’s HSD test (*P* < 0.05). **(b)** Alluvial diagram showing the relationships among *k-mean*s clusters, haplotype groups, continent of origin, row type, and growth type for the AGG accessions.

